# Anesthesia-triggered dopaminergic bursts actively induce forgetting: a paradigm shift in understanding cold-shock amnesia

**DOI:** 10.64898/2025.12.15.694449

**Authors:** Prachi Shah, Juan Carlos Martinez-Cervantes, Jacob A. Berry, Isaac Cervantes-Sandoval

## Abstract

Retrograde memory loss, the inability to recall events preceding amnesia onset, is a well-documented consequence of anesthesia. Recent discoveries, pioneered by *Drosophila* research, have challenged the view that forgetting is a passive process, and have demonstrated that rather it is a highly active, well-regulated biological process. This work builds on this and makes a remarkable and unexpected discovery that anesthesia itself triggers a robust, widespread burst of activity in dopaminergic neurons. Here, we report that both cold-shock and CO_2_ anesthesia elicits strong activation in PAM and PPL1 dopaminergic neuron populations. This calcium activity is accompanied by robust synaptic release of dopamine. Strikingly, pharmacological experiments show that this response is input-driven, as it is completely abolished by Na+ channel blockers and dampened by nAChR antagonists. Using behavioral methods, crucially, we show that anesthesia-induced amnesia can be prevented by blocking activity in PAM and PPL1 neurons during anesthesia. Together, our findings reveal a previously unrecognized active mechanism by which anesthesia induces forgetting, mediated by rapid dopaminergic signaling. This paradigm-shifting discovery redefines how we think about anesthesia-induced amnesia and the biology of forgetting.

## Main Text

Traditionally, studies in neuroscience of learning and memory have emphasized processes of acquisition and consolidation, but in the last decade, there has been increased efforts to understand the neurobiology underlying the opposing process – forgetting. Historically, forgetting has been debated on whether it occurs through a neuronal activity-dependent process or through passive mechanisms^1^. The psychological viewpoint of passive forgetting does not consider the brain as having the capacity to actively degrade the substrates of memory. However, we now have extensive evidence that forgetting is an active, well-regulated biological process that operates with dedicated pathways to cause memory loss^2–9^. Pioneering studies using *Drosophila* have allowed us to begin to understand the mechanism driving active forgetting. During olfactory memory acquisition, odors are assigned positive or negative value through reinforcement, produced by the coincident activation of a sparse set of Kenyon Cells (KCs) by an odorant and dopaminergic neurons (DANs) that innervate the 15 tile-like compartments of the mushroom body (MB) lobes ^10–16^. Each tile is linked to a specific mushroom body output neuron (MBON), whose activation drives either approach or avoidance behavior^14^. The pairing of an odor with a punishing or rewarding signal is thought to modify KC-to-MBON synaptic weights, allowing dopamine-induced plasticity to bias the MBON network toward the appropriate behavioral response^5,9,14,15,17–23^. Recent findings show that ongoing DAN activity, which reflects the animal’s behavioral state, also promotes memory forgetting^5–7,24,25^.

In addition to normal forgetting, retrograde amnesia—an inability to recall recent past events—can result from various insults, including head injury, stroke, or anesthesia. In *Drosophila*, cold-shock–induced anesthesia is commonly used after olfactory conditioning to examine the dynamics of memory consolidation. When flies are exposed to cold shock shortly after training, they exhibit graded retrograde amnesia^26^. Traditionally, this memory loss has been attributed to broad physiological disruptions caused by low temperatures – such as interference with metabolic and enzymatic processes, synaptic signaling, and other molecular processes. However, it remains unclear whether cold-shock anesthesia might engage specific neural circuits that promote forgetting, rather than simply causing passive biochemical shutdown. Here, we tested the hypothesis that anesthesia drives amnesia in a DAN activity-dependent manner.

### Burst of activity during anesthesia onset

To investigate whether anesthesia-induced amnesia is mediated by dopamine (DAN) signaling, we first characterized calcium responses in PPL1 and PAM DANs during cold-shock induced anesthesia. To do this, we performed functional imaging while inducing cold-shock anesthesia. We defined anesthesia onset as the abrupt cessation of locomotion, which occurred around ∼6°C and matched the anesthesia onset observed in freely moving animals. (Fig. 1a-b; Extended Data Fig.1). The anesthetic state was fully reversible, and flies rapidly regained locomotion once the temperature was restored, confirming a key property of anesthetics. In addition, we employed a physiological marker to confirm an anesthetized state by recording stimulus-evoked responses from neurons beyond the sensory layer. During anesthesia, the PPL1-α’3 DAN—which normally responds to odor stimulation even when flies are at rest—no longer showed any response to the odor (Extended Data Fig. 2), indicating that anesthesia disrupted sensory processing. Strikingly, we show that PPL1 and PAM dopamine neurons have massive bursts of calcium activity during cold-shock anesthesia (Fig. 1e, g). Several features of these responses stand out. **(1)** The calcium burst aligns to the transition from the awake to an anesthetized state (Fig 1b, d, Supplementary Video 1). **(2)** The magnitude is extremely large: most DANs have large responses to anesthesia, with some individual flies peaking ∼600 %ΔF/Fo, which is larger than typical responses to sensory stimuli. (Fig 1b, e, g). **(3)** Calcium responses peaked close in time to anesthesia onset, then partially decayed but remained elevated until the anesthetic was removed and the flies awoke (Fig 1b, e, g). **(4)** PPL1 and PAM neurons show distinct response profiles. PPL1 neurons begin increasing activity as the saline temperature drops (23-16°C, Fig.1i) and then show a sharp calcium surge at anesthesia onset; in contrast, PAM neurons show little response to initial temperature decrease (23-16°C, Fig.1m) and show a sharper surge at anesthesia onset (Fig. 1e, g, I, j, m, n). This agrees with previous findings that PPL1 but not PAM respond to mild cooling^27^. **(5)** Activation levels also vary within DAN subtypes. Notably, PPL1-γ1pedc exhibits a significantly smaller calcium response to cold-shock anesthesia than other PPL1 neurons, a difference confirmed using a split-GAL4 line specific to PPL1-γ1pedc (Fig. 1k and Extended Data Fig. 3). To further validate these results, we used the CRTC::GFP system^28^, which reports neural activity through activity-dependent translocation of CRTC from the cytoplasm to the nucleus. Cold-shock–anesthetized flies showed increased CRTC nuclear localization in both PPL1 and PAM-γ5 neurons compared to non-anesthetized controls (Extended Data Fig. 4). Together, these findings demonstrate that cold-shock anesthesia triggers large calcium responses in both PAM and PPL1 DANs.

**Fig. 1.**
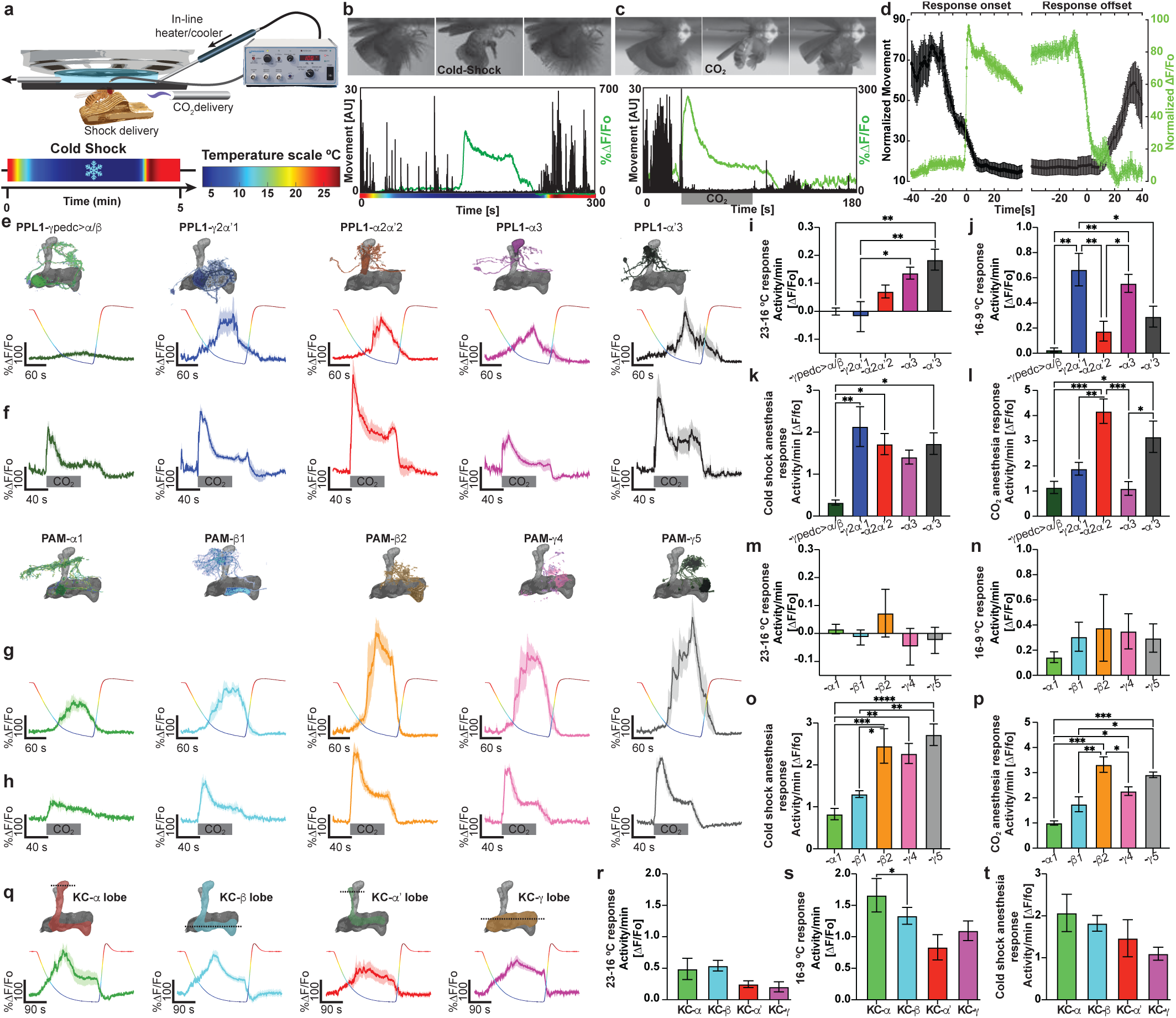
Dopaminergic neurons have robust calcium responses at the onset of anesthesia. **a,** Diagram of experimental set up to record in vivo anesthesia-induced responses under the microscope. Flies are mounted to a chamber where their body is free, but head if fixed with optical access to their brains. Cold-shock anesthesia is induced by decreasing the temperature from 25°C to 1°C using an in-line heater/cooler Peltier. CO_2_-anesthesia is induced by delivering the gas to the fly antenna through a Teflon tube. The temperature scale shows the change in temperature the fly experiences during the recordings. **b,** Example of DAN calcium response during cold shock-induced anesthesia, Upper panel, minimal projections of a single fly before, during and after cold-shock anesthesia. Bottom panel, movement of single fly quantified throughout cold-shock anesthesia aligned to GCaMP6f signal (green). **c,** Example of DAN calcium response during CO_2_-induced anesthesia. **d**, Alignment of multiple flies normalized movements and GCaMP6f recordings during the awake-anesthesia and anesthesia-awake transitions. The graph shows that the burst of activity occurs consistently at the onset of anesthesia. n=8. **e,** Calcium traces showing response profile of each PPL1 subtype during cold-shock anesthesia. Multicolored overlapping trace shows the temperature change. **f,** Calcium traces showing response profile of each PPL1 subtype during CO_2_-induced anesthesia. Gray bar indicates time of CO_2_ delivery. **g,** Calcium traces showing response profile of five PAM subtype during cold-shock anesthesia. Multicolored overlapping trace shows the temperature change. **h,** Calcium traces showing response profile of five PAM subtype during CO_2_-induced anesthesia. Gray bar indicates time of CO_2_ delivery. **i,** Average activity per minute of each PPL1 subtype during temperatures ranging from 23°C to 16°C. **j,** Average activity per minute of each PPL1 subtype during temperatures ranging from 16°C to 9°C. **k,** Average activity per minute of each PPL1 subtype during cold-shock anesthesia. **l,** Average activity per minute of each PPL1 during CO_2_ anesthesia. **m,** Average activity per minute of five PAM subtypes during temperatures ranging from 23°C to 16°C. **n,** Average activity per minute of five PAM subtypes during temperatures ranging from 16°C to 9°C. **o,** Average activity per minute of five PAM subtypes during cold-shock anesthesia. **p,** Average activity per minute of each PAM during CO_2_ anesthesia. **q,** Recordings of dopamine release using GrabDA showing response profile of each KC subtype during cold-shock anesthesia. Multicolored overlapping trace shows the temperature change the fly is experiencing. **r,** Average activity per minute of each KC subtype during temperatures ranging from 23°C to 16°C. **s,** Average activity per minute of each KC subtype during temperatures ranging from 16°C to 9°C. **t,** Average activity per minute of each KC subtype during cold-shock anesthesia. One-way Anova, followed by Tukey’s multiple comparisons, *p<0.05, **p<0.01, ***p<0.001, ****p<0.0001. Error bars indicate SEM.

To test whether the calcium bursts triggered neurotransmitter release, we monitored dopamine directly by expressing the GrabDA reporter in KCs. As with calcium, we observed a large dopamine surge at anesthesia onset (Fig. 1q-t).

We then questioned if this activity was a consequence of the temperature change or could it be generalized to other forms of anesthesia. As a result, we repeated the imaging experiment but recorded activity from PPL1 and PAM neurons while exposing the fly to CO_2_ under the microscope. Strikingly, we see massive bursts of activity when the fly is anesthetized by CO_2_ (Fig 1c, f, h and Supplementary Video 2). CO_2_ responses are similar to cold-shock responses, in that **(1)** the burst aligns the transition from the awake to an anesthetized state and **(2)** the calcium response in sustained and goes back to base level only after anesthetic is released (Fig. 1c). In contrast, CO_2_ responses differed in that the initial response to temperature change is no longer observed, so responses in both PPL1 and PAM have a sharper slope. Additionally, the difference between responses of different DANs seem to be partially maintained from cold-shock to CO_2_ anesthesia (i.e. PPL1-γ1pedc has overall smaller responses – Fig. 1 l, p). These results suggest that the burst of activity during anesthesia onset in not a physicochemical consequence of the temperature itself since other types of anesthesia induces similar responses even when the mechanism for anesthesia induction is completely different.

We then explored if anesthesia activated other memory relevant neurons. Our data shows all KC types and MBONγ2α’1 display an initial response as the temperature changes followed by DAN-like responses to the onset of cold-shock induced anesthesia (Extended Data Fig. 5). Dual calcium reporter imaging show that both DAN and KC responses are coordinated and occurred simultaneously (Supplementary Video 3). In contrast, when doing KC recordings during CO_2_ anesthesia, we strikingly found that KC responses were significantly delayed to anesthesia onset – on a scale of about 10 s. Finding that was further confirmed by dual calcium imaging (Supplementary Video 4). In addition, calcium responses to CO_2_ anesthesia in KCs appear as an initial decrease in calcium activity followed by a robust and sharp increase in activity (Extended Data Fig. 5). The distinct timing and response profiles across anesthetic types likely reflect fundamental differences in how cold-shock and CO_2_ alter neural activity during anesthesia, though the neural basis of the KC delay remains unknown. Together, these findings overturn the long-held view that the anesthetized brain is globally inactive, instead revealing widespread and robust activation across multiple neuron types.

### Cellular and circuit basis of anesthesia induced activity

We next sought to understand the circuit and cellular mechanisms underlying anesthesia-induced activity. Two main observations led us to suspect that the responses were input-driven rather than purely cell intrinsic mechanisms. First, the similarity between cold-shock and CO_2_-evoked responses indicated that they could not be explained solely by temperature-dependent biophysical effects dependent on cellular intrinsic mechanisms. Second, the responses were not driven by sensory input: surgical removal of the antennae—housing the primary thermos- and CO_2_-sensory neurons—left anesthesia-induced bursts intact, with only a small reduction in the initial cooling response (Extended Data Fig. 6). To begin dissecting the source of this activity, we tested whether it was driven by changes in general synaptic input, specifically whether a widespread loss of inhibitory tone during anesthesia could produce the observed excitatory burst across the network. To test this hypothesis, we recorded cold-shock responses before and after applying the GABAa receptor agonist Gaboxadol (1 mM), which should enhance inhibition and potentially rescue for a loss of inhibitory tone. After an initial cold-shock recording, we perfused Gaboxadol over the brain for 10 minutes and then repeated the cold-shock protocol. Contrary to our prediction, anesthesia-evoked responses were unchanged after drug application (Extended Data Fig. 7a-c). To further probe inhibitory contributions, we directly recorded from the GABAergic neuron MBON-γ1pedc>α/β during anesthesia and observed robust activity (Extended Data Fig. 7d). Together, these results indicate that enhancing GABAergic transmission does not alter the anesthesia-induced burst, suggesting that loss of inhibition is not its primary driver.

We next tested whether broadly suppressing neural communication would affect anesthesia-evoked calcium activity by blocking voltage-gated sodium channels with TTX. Applying 1 µM TTX—an established effective concentration—immediately altered the response profile. Before TTX, cold-shock produced the expected pattern: a gradual increase in activity during cooling, followed by a large burst at anesthesia onset (∼6 °C; ∼100 s after recording began). After TTX perfusion, the temperature-evoked activity was completely eliminated, and the anesthesia-induced burst was markedly delayed by ∼53 s (Fig. 2a-d). Interestingly, although TTX nearly abolished movement, a brief clonus-like leg twitch was still observed in some flies just before and during the delayed calcium rise, suggesting that both neural activity and the onset of anesthesia itself were delayed. Increasing TTX to 5 or 10 µM further delayed responses by ∼241 s and ∼343 s, respectively, and in some flies completely eliminated them (Fig. 2e-j). We hypothesized that incomplete drug penetration into the ventral nerve cord (VNC) might allow residual activity. To address this, we opened a small window in the thoracic cuticle to expose the VNC. Consistent with our hypothesis, recordings with both the brain and VNC exposed showed complete abolition of anesthesia-induced activity in the presence of TTX (Fig 2k-n). Surprisingly, our results indicate that anesthesia-induced activity is circuit-mediated and depends critically on input from the ventral nerve cord (VNC). Consistent with this, VNC severance markedly disrupted the characteristic anesthesia-evoked activity in DANs (Extended Data Fig. 8). These findings suggest that communication between the brain and VNC is required for cold-shock–induced responses—either because ascending pathways actively drive the burst or because reciprocal interactions between ascending and descending pathways are necessary for its initiation.

**Fig. 2.**
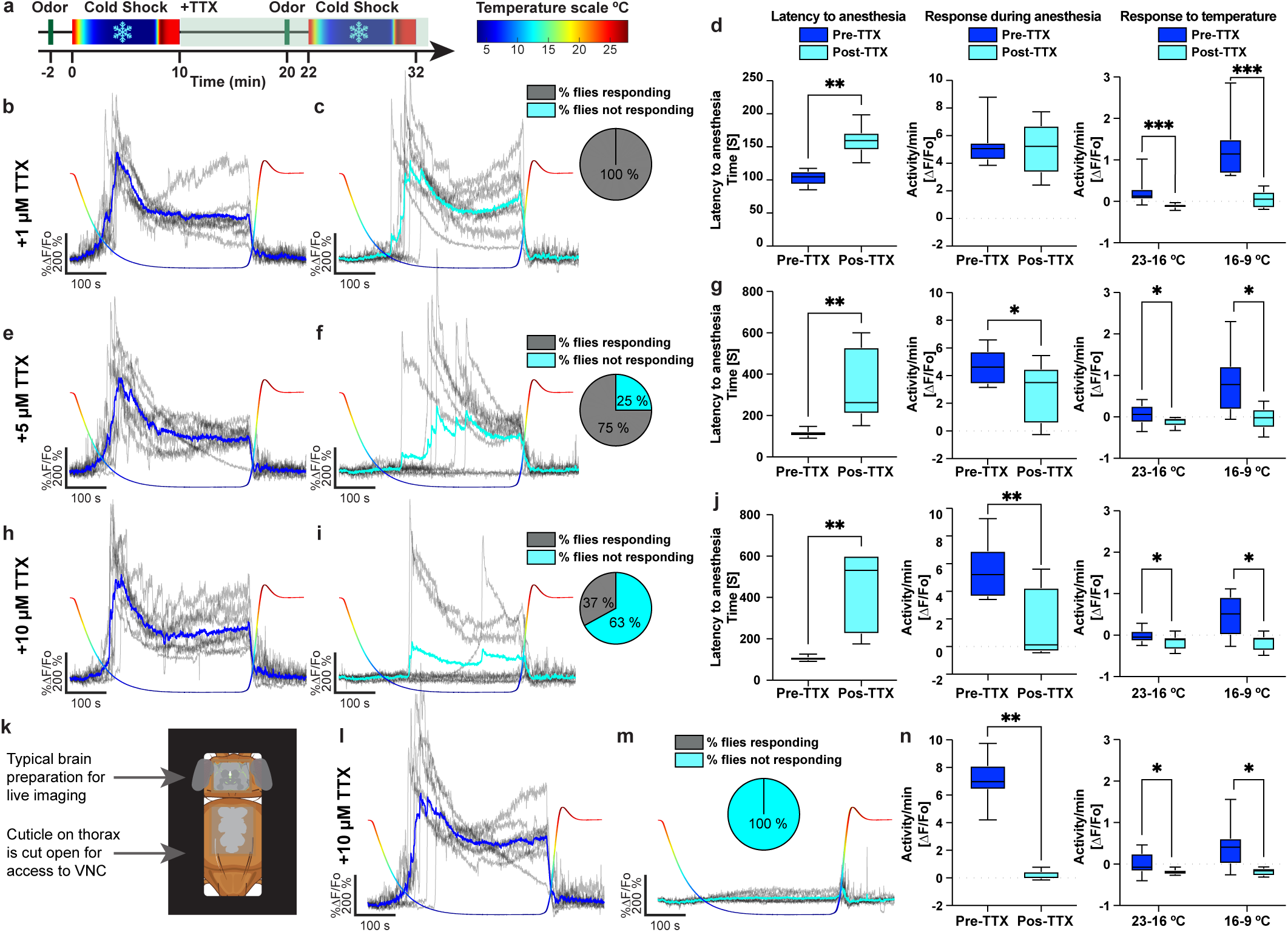
Tetrodotoxin delays latency to onset of anesthesia-induced response and VNC inhibition completely abolishes it. **a,** Diagram of experimental design. Each fly was imaged twice for 10 min. Baseline of calcium in PPL1-α2α’2 was recorded for 10 seconds before cold-shock. The temperature was increased back to 25°C at 7.5 min. Then, saline with different concentrations of TTX is perfused for 10 minutes. Cold-shock recording timeline is repeated post TTX treatment. Temperature recordings are shown as multicolor scale for each calcium trace. Scale is shown with corresponding colors. **b,e,h,l,** Cold-shock induced anesthesia responses in DAN pre TTX perfusion. Gray traces indicate individual fly responses; blue trace is the average anesthesia induced response. **c,** Cold-shock induced anesthesia responses in PPL1 post 1µM TTX perfusion. Gray traces indicate individual fly responses; blue trace is the average anesthesia induced response. Pie chart shows that 100% of flies displayed an anesthesia induced response post TTX treatment. **d,** left panel shows latency to onset of anesthesia response pre and post 1 µM TTX treatment. Middle panel, average activity per minute during anesthesia pre and post 1 µM TTX treatment. Right panel, calcium response to temperature during two ranges, from 23°C to 16°C and 16°C to 9°C pre and post 1µM TTX treatment. **f,** Cold-shock induced anesthesia responses in PPL1 post 5µM TTX perfusion. Gray traces indicate individual fly responses; blue trace is the average anesthesia induced response. Pie chart shows that 33% of flies displayed an anesthesia induced response post TTX treatment. **g,** left panel shows latency to onset of anesthesia response pre and post 5 µM TTX treatment. Middle panel, average activity per minute during anesthesia pre and post 5 µM TTX treatment. Right panel, calcium response to temperature during two ranges, from 23°C to 16°C and 16°C to 9°C pre and post 5µM TTX treatment. **i,** Cold-shock induced anesthesia responses in PPL1 post 10µM TTX perfusion. Gray traces indicate individual fly responses; blue trace is the average anesthesia induced response. Pie chart shows that 100% of flies displayed an anesthesia induced response post TTX treatment. **j,** left panel shows latency to onset of anesthesia response pre and post 10 µM TTX treatment. Middle panel, average activity per minute during anesthesia pre and post 10 µM TTX treatment. Right panel, calcium response to temperature during two ranges, from 23°C to 16°C and 16°C to 9°C pre and post 10µM TTX treatment. **k,** Illustration showing dissection of thoracic cuticle to allow drug penetration to the ventral nerve cord (VNC) **m,** Cold-shock induced anesthesia responses post 10 µM TTX perfusion with VNC access. Gray traces indicate individual fly responses; blue trace is the average anesthesia induced response. Pie chart shows that no flies displayed an anesthesia response post TTX treatment. **n,** Left panel, average activity per minute during anesthesia pre and post 10 µM TTX treatment with VNC access. Right panel, calcium response to temperature during two ranges, from 23°C to 16°C and 16°C to 9°C pre and post 10µM TTX treatment with VNC access. Non-parametric Wilcoxon paired test for pair comparisons. Two-way Anova, for temperature range comparisons followed by Tukey’s multiple comparisons. *p<0.05, **p<0.01, ***p<0.001, ****p<0.0001. Box represents Q1, median, and Q3. Whiskers represents min and max.

To test whether excitatory circuit mechanisms drive this response, we blocked nicotinic acetylcholine receptors with mecamylamine. Similar to TTX, 10 µM mecamylamine delayed anesthesia responses by ∼65 s in PPL1 neurons (Extended Data Fig. 9 b-d). Increasing concentration to 100 or 500 µM, eliminated temperature-evoked activity, and anesthesia responses were further delayed ∼322 s and ∼371 s, respectively, with some flies failing to respond altogether (Extended Data Fig.9 e-j). Unlike TTX, opening the thorax did not abolish the anesthesia-induced response completely (Extended Data Fig.9 k-m). In contrast, blocking glutamate receptors with philanthotoxin had no effect on cold-shock responses (Extended Data Fig. 10). These data suggest that acetylcholine signaling strongly modulates this activity, but blocking cholinergic transmission alone is insufficient to eliminate the anesthesia-evoked burst.

### Mushroom body compartment memory traces in anesthesia induced forgetting

We then explored what are the effects of cold shock on recently formed learning-induced plasticity observed in the different MB compartments, and if there is any correlation with the degree of responses in the corresponding DAN during anesthesia. Training the fly under the microscope induced robust plasticity in MBON-γ2α‘1 as previously reported^5^, defined by robust depression to the odor paired with shock (CS+) and significant potentiation to the unpaired odor (CS–). Cold-shock post-training completely restored odor responses to that of pre-training levels (Fig. 3a-e). In contrast, mock cold-shock did not show any restoration (Extended Data Fig. 11). Interestingly, and differing from MBONγ2α’1, plasticity in MBON-γ1pedc did not show a complete odor restoration to the CS+ (Fig. 3f-g). In contrast, CS– responses were completely restored after anesthesia which suggest that the CS– memory trace formed is weaker than that of the CS+. Finally, when we looked at a compartment previously implicated in the formation of LTM MBON-α3^16,20^, training once again induced robust plasticity in both CS+ and CS–. Notably, post-anesthesia response to the CS+ although partially restored it, is still significantly depressed compared to pre-training responses (Fig. 3h-i). Once again, CS– showed complete restoration. When we compared the level of recovery by calculating the degree of change from pre-training to post-anesthesia we observed that CS+ plasticity in MBON-γ1pedc and MBON-α3 are more resistant to anesthesia induced amnesia when compared to the MBONγ2α’1 compartment (Fig. 3j-k). We speculate that this data could be explained by the degree of PPL1 activation during cold-shock as this correlates to the degree of plasticity reversal. Across compartments, the anesthesia induced amnesia observed to the CS- is not significantly different between the different MBONs. This could be due to the weaker nature of the CS- memory and thus making it increasingly labile for anesthesia to disrupt it.

**Fig. 3.**
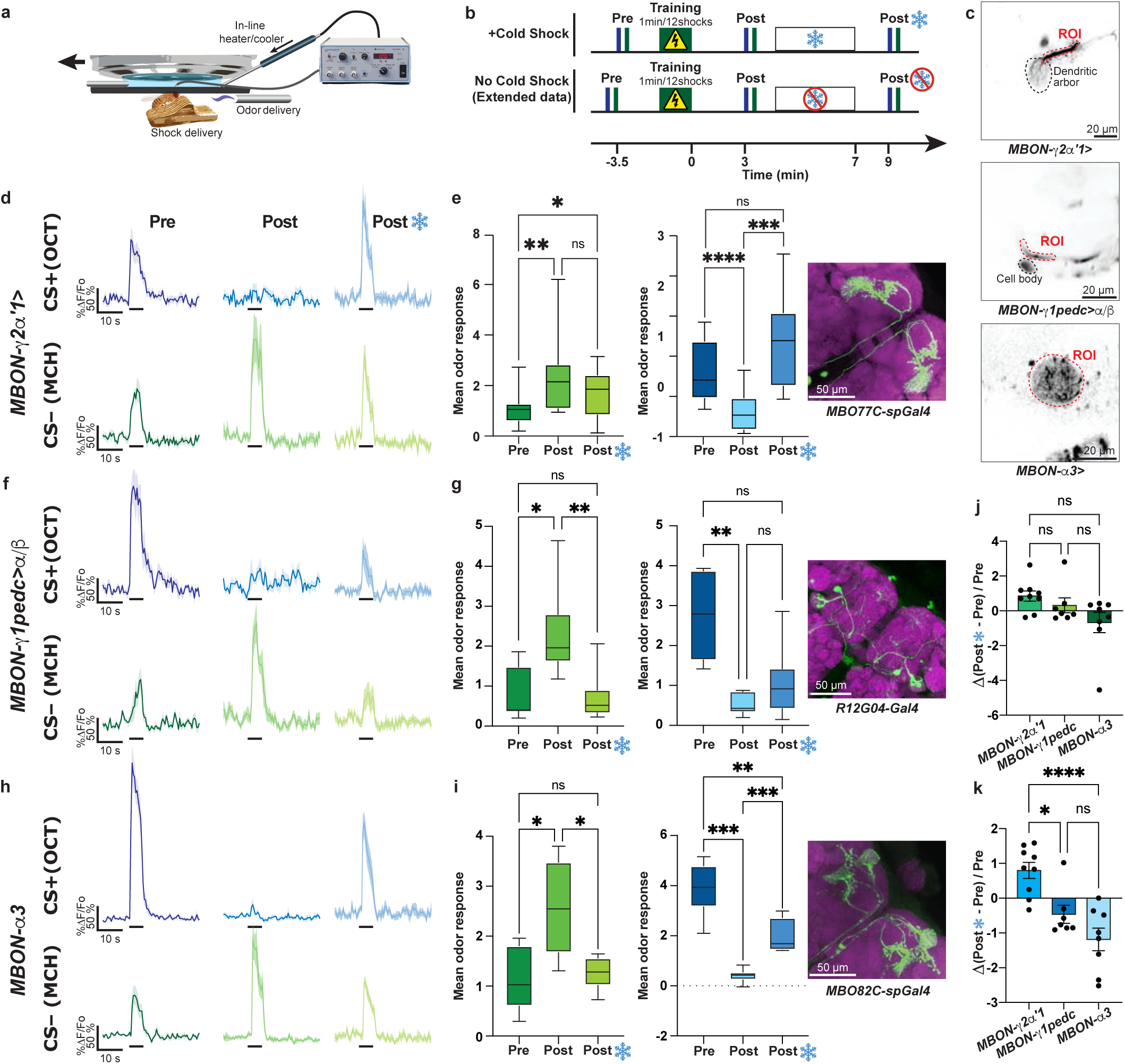
Anesthesia differentially affects distinct MBON memory traces. **a,** Diagram illustrating in vivo, under the microscope aversive olfactory conditioning training set up and an in-line heater/cooler to deliver cold-shock anesthesia delivery. **b,** Experimental design. Upper panel, Pre-training odor induced calcium response are recorded by presentation of 5-s 4-methylcyclohexanol (MCH) and 3-Octanol (OCT) with a 30-s ISI; 5 minutes later, flies were aversively trained by pairing 1 min presentation of OCT with 12-90 V shocks. Post-training odor induced calcium response were recorded 3 min after. Flies were then subjected to cold-shock anesthesia under the microscope. Two min after anesthesia recovery, post-cold shock anesthesia odor responses were recorded. Bottom panel, same as upper panel but flies are not anesthetized. **c,** Mean time series projection of baseline *GCaMP6f* signal in MBON-*γ*2*α*‘1 (upper), MBON-*γ*1pedc>*α*/*β* (middle), and MBON-*α*3 (lower) showing region of interest recorded. Scale bar: 20 µM. **d,f,h**, Odor response traces to CS+–OCT (top panel) and CS- –MCH (bottom panel) for pre-training, post-training and post-cold shock anesthesia for MBON-*γ*2*α*‘1 (d), MBON-*γ*1pedc>*α*/*β* (f), and MBON-*α*3 (h). Thick lines represent time of odor presentation. **e,** MBON-*γ*2*α*‘1 MCH (CS-) responses were significantly potentiated post-training and anesthesia compared to pre-training. OCT (CS+) responses were significantly suppressed post-training, nevertheless, the plasticity was completely reversed following cold-shock anesthesia. **g,** MBON-*γ*1pedc>*α*/*β* MCH responses were is significantly potentiated post training, but not significantly different post-anesthesia when compared to pre-training. OCT (CS+) responses were significantly suppressed post-training. In contrast, no significant differences were observed between post-training and post-anesthesia responses. **i,** MBON-*α*3 MCH (CS-) responses were significantly potentiated post-training, but not significantly different post-anesthesia compared to pre-training. OCT (CS+) responses were significantly suppressed post-training and post-anesthesia. **j,** No significant differences between each MBON to change in CS- responses post-anesthesia relative to pre-training. **k,** MBON-*γ*1pedc>*α*/*β*, and MBON-*α*3 show significantly less recovery in the response to the CS+ following anesthesia. One-way Anova followed by Tukey’s multiple comparisons test, *p<0.05, **p<0.01, ***p<0.001, ****p<0.0001. Box represents Q1, median, and Q3. Whiskers represents min and max. n=7-12.

### Blocking PPL1 and PAM dopamine activity during anesthesia inhibits forgetting

Our next goal was to test whether PPL1 and PAM DAN activity during anesthesia drives active forgetting. Although DANs are known to bidirectionally regulate learning and forgetting, our data show that anesthesia also strongly activates other memory-relevant neurons, including KCs and MBONs (Extended Data Fig. 5), raising the possibility that these cells contribute to anesthesia-induced amnesia. However, artificial activation using TrpA1 of KCs (*ok107-gal4*) or MBON-γ2α′1 for 20 min had no effect on memory performance (Extended Data Fig. 13b, c). In contrast, activating PPL1 neurons completely abolished memory, supporting a specific role for DANs in this process (Extended Data Fig. 13c). These experiments further support dopamine’s central role on active forgetting. However, several caveats remain. (1) Although, we show that a 10 S cold-shock is sufficient to cause memory loss (Extended Data Fig. 12a, b), it remains to be determined if longer artificial TrpA1 activation of KC or MBON reveals memory effects. (2) Because KCs provide major input to DANs ^29,30^, KC activation should indirectly activate dopamine neurons, yet we saw no memory change, an unexpected result. (3) We tested only one MBON type; broader MBON activation might yield different outcomes.

To test whether cold-shock amnesia is dopamine-dependent, we blocked both PPL1 and PAM DANs during anesthesia, as aversive-conditioning plasticity spans domains innervated by both types ^23,31^. Inhibition of DANs with the red-shifted anion channelrhodopsin RubyACR during cold shock significantly improved memory, although RubyACR expression itself modestly impaired learning (Fig. 4a-b). We therefore validated the results using *Shibire^ts^* to block dopamine release during CO₂ anesthesia, which also activates DANs and produces amnesia (Extended Data Fig. 12c, d). Again, memory was significantly preserved relative to genetic and temperature controls (Fig. 4c-d).

**Fig. 4.**
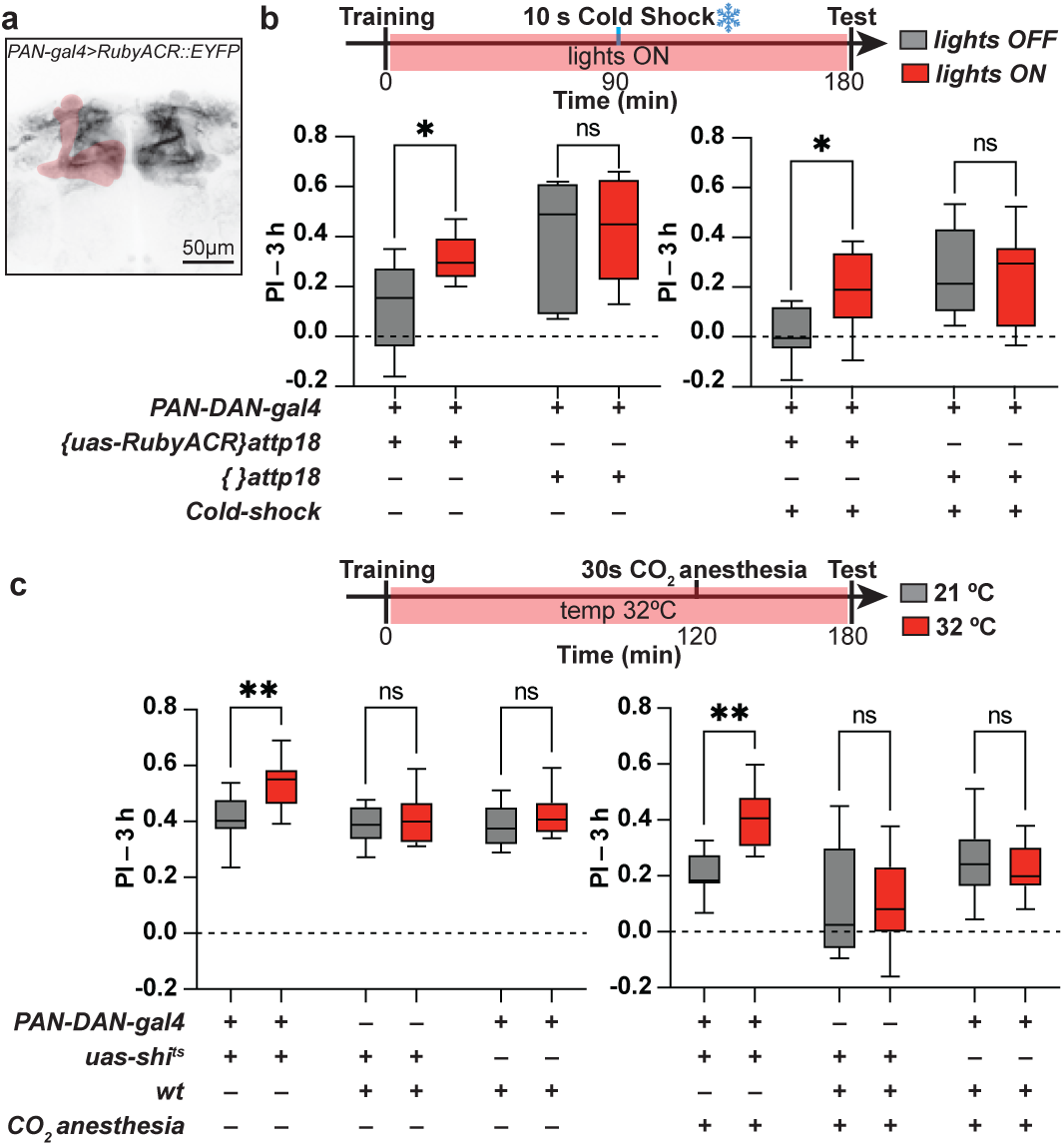
Blocking PPL1 and PAM dopamine activity during anesthesia is sufficient to inhibit forgetting. **a,** Expression pattern of TH-gal4/R58E02-gal4 (PAN-DAN-gal4) recombination line used to label both PPL1 and PAM neurons expressing RubyACR tagged with EYFP. **b,** Upper panel, diagram of experimental protocol. Flies were trained using single cycle aversive olfactory conditioning and placed in lights off or on conditions to inhibit DAN activity by RubyACR. Flies were cold-shock anesthetized for 10 seconds 90 minutes post training. Flies were tested for memory performance 180 minutes pos training. Left-lower panel, Memory is enhanced in flies where DAN activity is blocked post training compared to genetic and light controls. Right-lower panel, Memory is enhanced in flies where dopamine activity is blocked during cold-shock induced anesthesia compared to genetic and light controls. **c,** Upper panel, Diagram of experimental protocol. Flies were trained using single cycle aversive olfactory conditioning and placed at either 21°C or 32°C. Flies were given 30 seconds of CO_2_ anesthesia at 120 minutes post training and tested for their memory performance at 180 minutes post training, Left-lower panel, Performance was significantly enhanced when blocking PPL1 and PAM neurons post training with Shibire^ts^ compared to genetic and temperature controls. Right-lower panel, Memory was significantly enhanced in flies when dopamine release is blocked during CO_2_-induced anesthesia compared to genetic and light controls. Two-way Anova followed by Fisher’s multiple comparisons, *p<0.05, **p<0.01. Box represents Q1, median, and Q3. Whiskers represents min and max. n=8-10.

These findings strongly support that anesthesia-induced forgetting is an active, dopamine-driven process: blocking dopamine during anesthesia is sufficient to protect recently formed memories. More broadly, our results challenge the long-standing view that cold-shock anesthesia reflects passive biochemical shutdown and instead reveal a circuit-level mechanism in which anesthesia triggers widespread neural activation, with dopamine specifically driving retrograde memory loss.

## Discussion

Historically, anesthesia has been viewed as simply “turning off” the brain. However, accumulating evidence shows that anesthetics induce characteristic shifts in neural oscillations that disrupt communication between brain regions and produce unconsciousness—a behavioral state distinct from sleep^32^. EEG recordings demonstrate a transition from low-amplitude, fast oscillations to high-amplitude, slow-wave activity as patients lose consciousness^33,34^ reflecting large-scale changes in coordinated neuronal activity. Here, we provide the first demonstration of anesthesia-evoked large-scale neural activation in the *Drosophila* brain. We identify several striking features of this phenomenon. (1) Anesthesia induces robust calcium transients across multiple neuronal cell types. (2) These responses occur precisely at the onset of the anesthetic state. (3) Mechanistically distinct anesthetics elicit similar large-scale activation. (4) This activity is driven by synaptic input and intact communication between the brain and VNC. (5) Pharmacological delay of this activity suggests, although not conclusively, to also delay the onset of anesthetic state. (6) Dopaminergic activation during this event drives active forgetting of recently formed memories.

Our findings parallel a well-known neurophysiological event in vertebrates called spreading depolarization (SD), a wave of sustained neuronal depolarization accompanied by loss of excitability. SD was first described in 1944 in the context of spreading suppression of cortical activity during experimental epilepsy^35^. Although extensively studied, the mechanisms underlying SD remain difficult to probe due to the complexity of mammalian circuits. Our preparation provides a tractable model for studying analogous dynamics: the widespread depolarization observed at anesthesia onset resembles a spreading depolarizing event that transiently disrupts synaptic communication.

We also show that blocking dopamine activity during anesthesia prevents the retrograde amnesia typically produced by cold shock. This supports a conserved role for dopamine in active forgetting of labile memories and further indicates that memory degradation relies on dedicated biological mechanisms rather than passive erasure. Together, our results reshape the longstanding view that cold-shock amnesia simply reflects biochemical inactivation. Instead, we reveal an active, circuit-level process in which anesthesia triggers widespread neural activation, with dopamine signaling specifically driving memory loss. This work not only advances our understanding of forgetting mechanisms but also provides a framework for studying how major neural insults, such as seizures, traumatic brain injury, or intoxication, interact with memory systems.

## Methods

### Drosophila husbandry

Flies were cultured on standard medium at room temperature. Crosses, unless otherwise stated, were kept at 25 °C and 70% relative humidity with a 12-hr light-dark cycle. The following lines were used for experiments, crosses, and to generate stocks: *Canton-S*, *MB082C-gal4^14^, MB077C-gal4^14^*, *ok107*-*gal4*^36^*, R58E02-gal4^37^, MB315C-gal4^13^,; uas-shibire^ts^;, uas-gcamp6f^38^*, *uas-tdtomato^39^, uas-HfACR1::EYFP;;^40^*, *;uas-mCD8::mCherry-T2A-lacZ::nls, uas-CRTC::GFP;^28^*, MB-RGECO;MB-RGECO^41^, *{}attp18*, *{}attp40*.

### Immunostaining

Whole brains were isolated and processed with minor modifications of those described ^37^. Brains were first incubated with primary antibodies including: rabbit polyclonal anti-GFP (1:1,000, Life Technologies, cat# A11122), mouse monoclonal anti-nc82 (1:50, University of Iowa, DSHB), mouse anti-LacZ (1:500, Promega, cat#Z3781), chicken anti-mCherry (1:500, Aves, cat#ab_1910557). Secondary antibodies included: anti-rabbit IgG conjugated to Alexa Fluor 488 (1:800, Life Technologies Cat# A11008, RRID: AB_143165), anti-mouse IgG conjugated to Alexa Fluor 633 (1:1000, Life Technologies Cat# A21052, RRID: AB_141459), anti-chicken IgY conjugated to CF633 (1:1000, Biotum, cat#20104). Images were collected using a 20X objective with a Leica TCS SP8 confocal microscope with 488 and 633 nm laser excitation.

### In vivo imaging

Functional imaging was performed as previously reported ^17,42^. Briefly, a singly fly was gently aspirated without anesthesia into a metal pipette connected to the vacuum (MicroGroup Hypodermic Tubing 304H22) to immobilize the head using proboscis aspiration. Once the head is immobilized, using a micromanipulator, the fly was inserted in a narrow slot the width of their body in a custom-designed recording chamber. The head was then fixed by gluing the sides of the eyes and thorax to the chamber using melted myristic acid. After this the fly was released from the metal pipette and the proboscis is fixed with myristic acid to avoid brain movement during proboscis extension. Using a syringe needle (BD PrecisionGlide^TM^ Needle REF 305193), a small, square section of dorsal cuticle was removed from the head to allow optical access to the brain. Fresh insect Ringer’s solution (103 mM NaCl, 3 mM KCl, 5 mM HEPES, 1.5 mM CaCl_2_, MgCl_2_, 26 mM NaHCO_3_, 1 mM NaH_2_PO_4_, 10 mM trehalose, 7 mM sucrose, and 10 mM glucose [pH 7.2]) was perfused immediately across the brain to prevent desiccation and ensure the health of the fly. Then the fat bodies and trachea above the brain was removed. Using a 20x water-immersion objective and a Leica TCS SP8 II confocal microscope with a 488 nm solid state laser, we imaged MBONs for 2 min at 2 Hz, during which odor stimuli was delivered starting 30 seconds after imaging initiation. We used one HyD channel (510-550 nm) to detect GCaMP6f fluorescence and 600-785nm to detect Tdtom. Recordings were performed at 2 Hz.

For DANs and KCs, fluorescence was acquired from a region of interest (ROI) drawn around the axons of the neurons of interest. For MBON-γ2α’1, γ1<pedc, and α3, fluorescence was acquired from a region of interest (ROI) drawn around the dendrites of the neurons of interest. ΔF/Fo was calculated using a custom MATLAB script.

### Anesthesia under the microscope

To induce cold-shock anesthesia under the microscope, saline solution was perfused at 2 ml/min throughout the imaging. The temperature of the saline was modulated with an inline heater/cooler (Warner Instruments), monitored with a thermistor placed 2 mm lateral and 1 mm anterior to the fly’s head, and recorded via an analog-to-digital converter using an Arduino microcontroller connected to a custom-made Matlab data acquisition script. Temperature data was collected at 72 Hz. Neuronal activity was first recorded for 10 s at 25 °C, after which the saline temperature was lowered to 0 °C in the instrument. Due to heat exchange, the thermistor typically registered a minimum of ∼3.5 °C. The temperature was held at 0 °C for 4 minutes and then returned to 25 °C. Flies typically entered in an anesthetic state at around ∼6 °C. Regions of interest were drawn around neuropil of dopaminergic neurons innervating the mushroom body and the mean fluorescence intensity was calculated. Each fly was imaged up to three to record responses of different DANs. Flies were allowed to recover for 5 min between each recording. A separate set of animals was recorded presenting only one stimulus to ensure that its presentation order did not affect the experiment results, i.e., there was no significant difference in the response amplitudes between brains when the cooling stimulus was presented first versus last.

For CO_2_ anesthesia under the microscope, flies were dissected as indicated above and CO_2_ was delivered by switching a clear air stream to a CO_2_ stream using solenoids valves (The Lee company, cat# LFRA1220170D). Both, the air and CO_2_ streams were maintained at 1000 ml/min and routed through a Teflon tubing (∼2.5 mm diameter) to the fly. Air and gas pressures were regulated using Alicat Mass Flow Controllers (Alicat 500CC, cat# BC-500C-N2V25A and Alicat 2L, cat# BC-002L-N2V25A). The solenoids controlling the CO_2_ delivery was controlled by an Arduino microcontroller (Arduino Uno) with custom-made programs.

#### Drug administration experiments

For pharmacological manipulations, we first recorded a baseline cold-shock anesthesia response as described above. Immediately afterward, saline containing the drug of interest was perfused continuously for 10 minutes to ensure robust exposure of the brain tissue. Following this incubation period, a second cold-shock response was recorded using the same temperature protocol. This pre-/post-drug design allowed us to directly compare how each compound altered the dynamics of the anesthesia-evoked activity within the same animal.

#### Thoracic cuticle removal

In a subset of experiments, the thoracic cuticle was removed in a manner similar to the head capsule dissection to improve drug access to the ventral nerve cord (VNC). During these experiments, we ensured that the fly’s orientation allowed the perfused saline to directly bathe the thoracic opening throughout the imaging session, maximizing drug penetration into the VNC.

#### VNC severance surgery

For VNC severance experiments, an initial cold-shock response was recorded as a baseline. The fly was then removed from the stage, and the VNC was surgically severed using a fine syringe needle (BD PrecisionGlide™ Needle, REF 305193). Care was taken to minimize collateral damage to surrounding tissues. After the lesion, the fly was repositioned under the microscope, allowed to stabilize, and imaged again while undergoing a second cold-shock stimulus. This approach enabled us to directly assess how loss of ascending and descending VNC–brain communication altered the anesthesia-induced activity.

### CRTC imaging

To assess neuronal activation following cold-shock using the CRTC^28^ system, individual flies expressing CRTC components in PPL1 and PAM neurons were transferred directly from their food vials into pre-chilled glass vials. These vials had been cooled in an ice bucket for at least 30 minutes prior to the experiment to ensure stable low temperatures. Flies remained in the cold vial for 2 minutes, after which they were decapitated immediately. Control flies were decapitated immediately without anesthesia by immobilizing them in a fly aspirator^42^. Brains were then dissected and processed for immunohistochemistry, staining for CRTC::GFP, LacZ::nls, and mCherry::CD8 as described above. Neuronal activation was quantified by measuring the nuclear localization index (NLI) of CRTC::GFP, following previously published procedures. Briefly, for each neuron we calculated the NLI as:

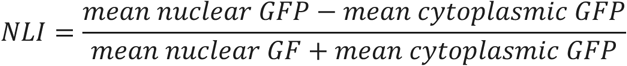

This index ranges from −1 to +1 and reflects the relative enrichment of CRTC::GFP in the nucleus versus the cytoplasm for individual neurons, thereby providing a quantitative readout of CRTC activation following cold-shock.^3030^

### Odor presentation

To deliver odors to flies under the microscope, a stream of air (100 ml/min) was diverted (via solenoids (The Lee company, cat# LFRA1220170D) from flowing through a clean 20 ml glass vial to instead flow through a 20 ml glass vial containing a 0.5 µl drop of pure odorant. This air stream was then serial diluted into a larger air stream (1500 ml/min) before traveling through Teflon tubing (∼2.5 mm diameter) to reach the fly. The air pressure for the odor stream and main large air stream were regulated using Alicat Mass Flow Controllers (Alicat 500CC, cat# BC-500C-N2V25A and Alicat 2L, cat# BC-002L-N2V25A). Both, solenoids controlling odor delivery and the Grass stimulator that delivers shocks were controlled by Arduino microcontroller (Arduino Uno) with custom-made programs (available upon request). The regular training protocol consisted of one 60 s odor presentation of 3-octanol (OCT– Sigma, cat#218405), simultaneously paired with 12, 90 V electric shocks. The pre and post training odor responses were collected by presenting a 5 s presentation of 4-methylcyclohexanol (MCH– Sigma, cat#589-91-3) followed by a 30 second rest of clean air and then another 5 s presentation of OCT.

For odor responses quantification a baseline was calculated as the mean fluorescence across the 5 s before each odor presentation. This baseline was then used to calculate %Δ*F*/*F*_o_ for the complete recording. Bar graphs represent distribution of %Δ*F*/*F*_o_ responses across the 5 s of odor presentation. Solid lines in fluorescence traces represent mean %Δ*F*/*F*_o_ ± standard error (SE) (shaded area) across the odor responses.

### Fly movement analysis

A camera (Fig.1a, FLIR Grasshopper3 USB3.0) and lens (MVL7000 macro lens, Navitar) was used to record the fly behavior under the confocal while simultaneously recording Ca^2+^ activity in DANs. In order to quantity physical movements (body movement) simultaneously with DAN activity during *in vivo* imaging experiments, the video camera was triggered using an Arduino microcontroller to capture a frames simultaneously with confocal frame scan. Videos of fly locomotion were captured at 60 Hz. For Fly motion quantification, raw AVI files were processed using custom ImageJ macros that first utilize the “Delta F down” plugin from the WCIF-ImageJ collection to perform image subtraction and then this result was thresholded (changes in pixel intensity between 5 and 255 then became 255). A circular region of interest (ROI) was centered on the fly body, and the integrated density of this ROI, divided by 255, was calculated to obtain the total number of pixels shift from frame to the next as a measure of fly motion over time.

### Behavior

#### Cold-Shock in freely moving flies

We designed a square behavioral arena consisting of a 20 × 20 mm Peltier element surrounded by a 3D-printed frame and covered with a thin, transparent acrylic lid. The arena was kept at 25 °C while flies were introduced. Individual flies were aspirated into the chamber and allowed to freely explore for 10 s before the temperature was lowered. The Peltier surface temperature was then decreased from 20 °C to 0 °C in 2 °C increments. Temperature control was achieved using an Arduino microcontroller operating the Peltier element through pulse-width modulation.

Behavior was recorded with a FLIR camera positioned above the arena. Fly trajectories were extracted and analyzed using FlyTracker (Eyjolfsdottir et al., 2014). From these trajectories, instantaneous walking speed was calculated for each fly and plotted as a function of time to quantify the behavioral effects of cold-shock.

#### Olfactory Conditioning

Groups of 50-60 flies were first equilibrated for ∼15 min in a fresh food vial to the environment of a behavioral room dimly lit with red light at 23°C and 70% humidity. They were then loaded into a training tube where they received the following sequence of stimuli: 30 sec of air, 1 min of an odor paired with 12 pulses of 90V electric shock (CS+), 30 sec of air, 1 min of a second odor with no electric shock (CS-), and finally 30 sec of air. For conditioning odors, we bubbled fresh air through 3-octanol (OCT) and 4-methylcyclohexonal (MCH) at concentrations of 0.1% and 0.05% in mineral oil, respectively. To test memory retention, we tapped the flies after conditioning back into a food vial, unless otherwise stated, for testing at a later time point as indicated in the figures. To measure the memory, flies were transferred into a T-maze where they were allowed 2 min to choose between an arm with the CS+ odor and an arm with the CS- odor. For all experiments, two groups were trained and tested simultaneously. One group was trained with OCT as the conditioned stimulus paired with reinforcer (CS+) and MCH unpaired with reinforcer (CS-), while the other group was trained with MCH as CS+ and OCT as CS-. Each group (50-60 flies) tested provided a half performance index (half PI): Half PI = ((# flies in CS- arm) – (# flies in CS+ arm)) / (# flies in both arms). A final PI was calculated by averaging the two half PI’s. Since the two groups were trained to opposite CS+/CS- odor pairs, this method balances out naïve odor biases.

To assess the effects of cold-shock anesthesia on memory performance, flies were transferred from their food vials into pre-chilled glass vials. These vials were cooled by placing them in an ice bucket for at least 30 minutes prior to the experiment. Flies remained in the cold vial for either 2 minutes, 1 minute, or 10 s, depending on the experimental condition. After cold-shock, flies were returned to food vials and allowed to recover before 3-hour memory testing.

For experiments involving RubyACR, flies were trained as described above but under extremely dim purple light to minimize unintentional optogenetic activation. After training, flies were transferred to food vials and placed on top of a red LED platform. The LEDs were pulsed at 15 Hz with a 30 ms pulse width and controlled by an Arduino microcontroller.

During cold-shock anesthesia, flies were transferred to a pre-chilled glass vial as described above; however, for optogenetic trials, the outside of the vial was surrounded by pulsing red LEDs to activate RubyACR during anesthesia. Following cold-shock, optogenetic flies were tested in the T-maze in complete darkness to avoid further light stimulation.

For *Shibire^ts^* experiments, flies were placed in a 32 °C incubator immediately after training. A flexible 1 mm inner-diameter silicone tube was inserted into each fly vial to allow CO_2_ delivery without removing the vials from the incubator. CO_2_ was injected through the tubing for 30 seconds to induce anesthesia, after which flies were allowed to recover at 32 °C until memory testing.

For *TrpA1* experiments flies were just moved in and out of a 32 °C incubator at the indicated times.

### Quantification and Statical Analysis

All replicates in the manuscript were biological replicates. Statistics were performed using Prism 9 (GraphPad). All tests were two tailed and significance levels were set at α=0.05. The figure legends present the *p* values and comparisons made for each experiment. Unless otherwise stated, non-parametric tests were used for all imaging data.

## Acknowledgements

This work was supported by grants R01GM147917 to IC-S; F31GM157963 from the NIGMS, T32AG071745 from NIA, and ARCS fellowships to PS. We acknowledge Glenn Turner from Janelia Farms for sharing RubyACR fly stocks. Finally, we acknowledge the MakerHub at Georgetown for support building custom made tools for this project.

---

**Extended Data Fig. 1.**
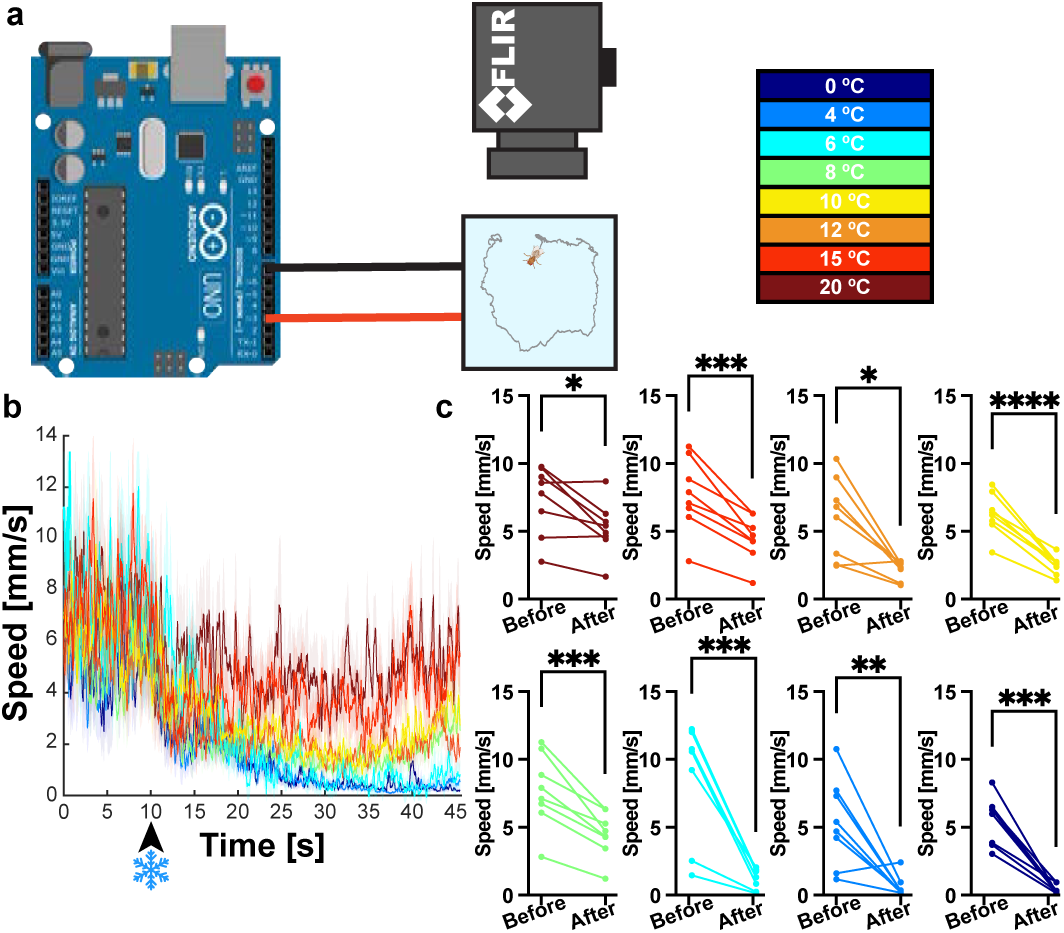
Freely moving *Drosophila* locomotion responses to temperature decreases. **a,** Behavior set up using an 20 x 20 mm Peltier thermoelectric cooler module controlled by an Arduino surrounded by a 3D-printed frame and covered with a thin, transparent acrylic lid. The arena was kept at 25 °C while flies were introduced. Individual flies were aspirated into the chamber and allowed to freely explore for 10 s before the temperature was lowered. Flies were subjected to different temperatures on the indicated in the color map. Temperatures were precisely maintained using an Arduino microcontroller. **b,** Behavioral traces of average fly speed over time at each tested temperature. **c,** Significant difference in average speed is observed for all tempera-tures tested before and during a cold-shock. At temperatures below 6°C, flies were fully anesthetized. Parametric paired t-test, *\*p<0.05, **p<0.01, ***p<0.001, ****p<0.0001. n=8*.

**Extended Data Fig. 2.**
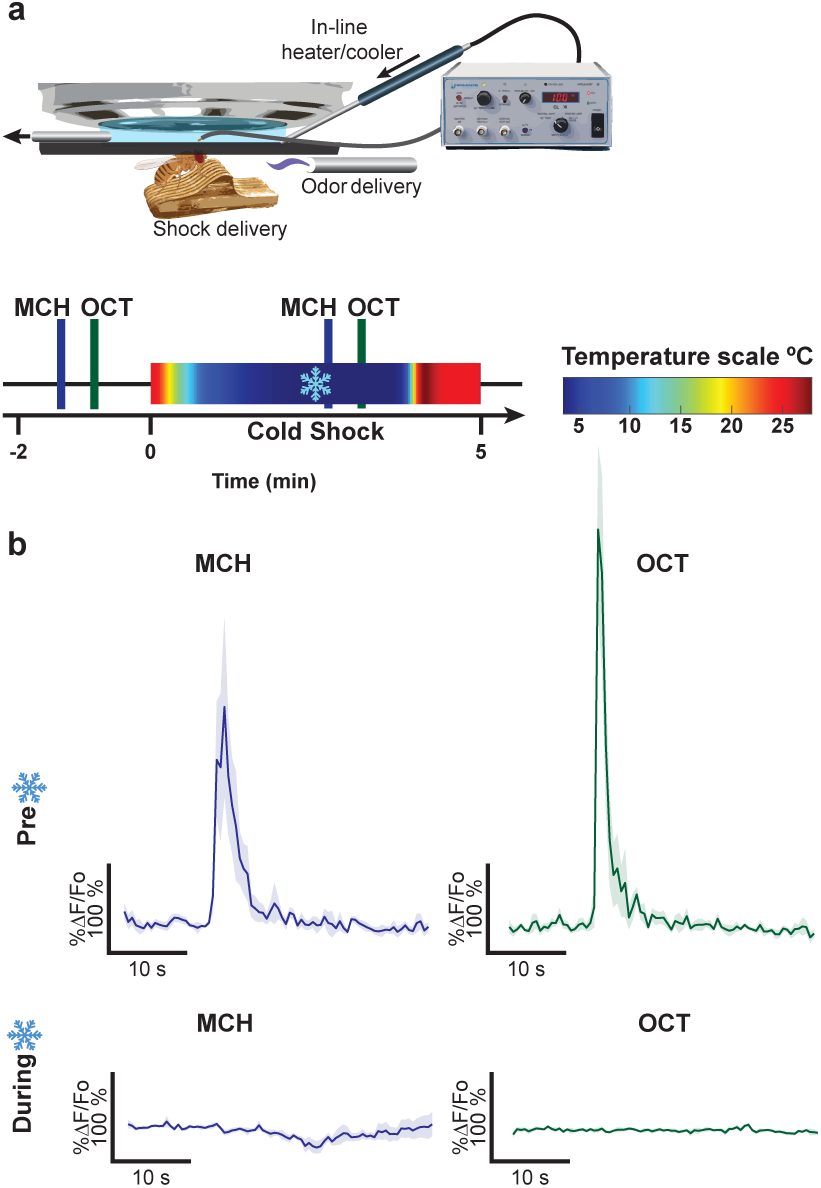
Anesthesia under the microscope suppresses olfactory responses in PPL1-α’3. **a,** Diagram of experimental set up to record in vivo anesthesia-induced responses under the microscope. Flies are mounted to a chamber where their body is free, but head is fixed with optical access to their brains. Cold-shock anesthesia is induced by decreasing the temperature from 25°C to 0°C using an in-line heater/cooler Peltier. The temperature scale shows the change in temperature the fly experiences during the recordings. Flies were exposed to 5 s odor pulses using MCH and OCT before experiencing cold-shock anesthesia. Flies were exposed to the same odors while anesthetized. **b,** Pre-cold-shock anesthesia odor-induced calcium responses are robust. During anesthesia, calcium responses to both MCH and OCT are completely abolished.

**Extended Data Fig. 3.**
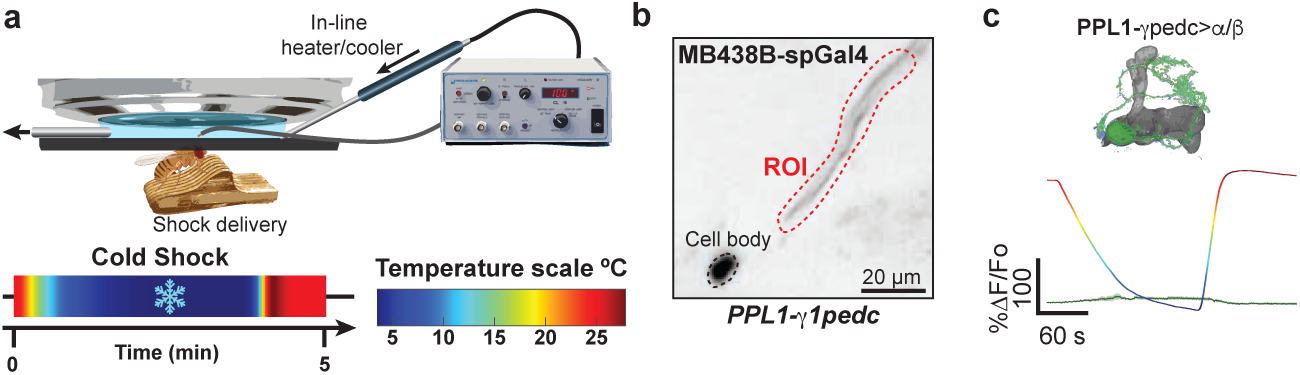
Calcium responses to Cold-Shcok anesthesia in PPL1-g1pedc>a/b using a the split-gal4 driver MB438B. **a,** Diagram of experimental set up to record in vivo anesthe-sia-induced responses under the microscope. Flies are mounted to a chamber where their body is free, but head is fixed with optical access to their brains. Cold-shock anesthesia is induced by decreasing the temperature from 25°C to 0°C using an in-line heater/cooler Peltier. The temperature scale shows the change in temperature the fly experiences during the recordings. **b,** Mean time series projection of baseline GCaMP6f driven by MB438B-gal4 in the PPL1**-**g1pedc>a/b projections and cell body. ROI is indicated by dotted red line. **c,** Calcium trace showing response profile of PPL1**-**g1pedc>a/b during cold-shock anesthesia. Multicolored overlapping trace shows the temperature change.

**Extended Data Fig. 4.**
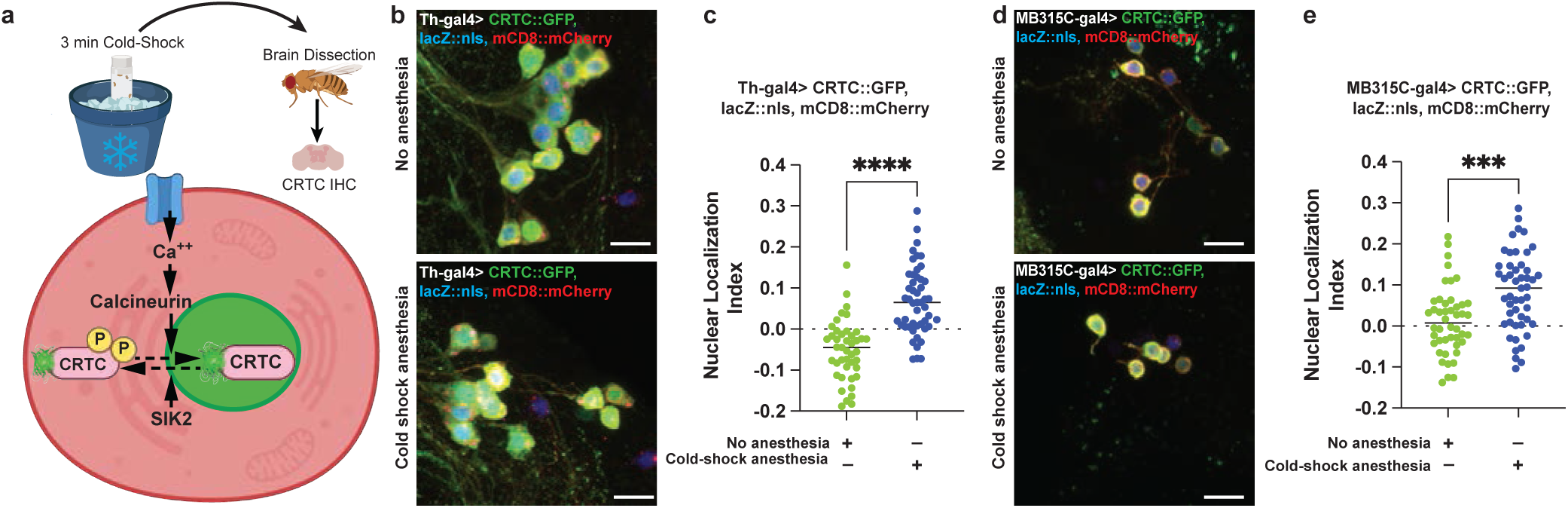
PPL1 and PAM dopaminergic neurons response to cold-shock anesthesia evaluated by CRTC nuclear translocation. **a,** Upper panel: experimental set up showing flies were cold-shock anesthetized and brains dissected and processed to be stained immediately after. Bottom panel: Schematic showing CRTC system. Activity-dependent translocation of CREB-regulated transcriptional co-activator (CRTC) from the cytoplasm to the nucleus is achieved by dephosphorylation of CRTC::GFP by calcium-dependent signaling. **b,** Representativ confocal images showing PPL1 cell bodies expressing CRTC::GFP using the TH-gal4. Upper panel, PPL1 cell bodies of control flies experiencing no cold-shock anesthesia. Bottom panel, PPL cell bodies CRTC signal after cold-shock anesthesia. **c,** The CRTC nuclear localization index is significantly increased in PPL1 neurons following cold-shock anesthesia when compared to contro group. **d,** Representative confocal images showing PAM-g5 cell bodies expressing CRTC::GFP using the MB315C-gal4. Upper panel, PAM-g5 cell bodies of control flies experiencing n cold-shock anesthesia. Bottom panel, PAM-g5 cell bodies CRTC signal after cold-shock anesthesia. **e,** The CRTC nuclear localization index is significantly increased in PAM-g5 neurons followin cold-shock anesthesia when compared to control group. Green labels, CRTC::GFP; blue labels, LacZ::nls for the nuclei; red labels, membrane mCD8::mCherry. Non-parametric Mann-Whitne test, ***p<0.001, ****p<0.0001.

**Extended Data Fig. 5.**
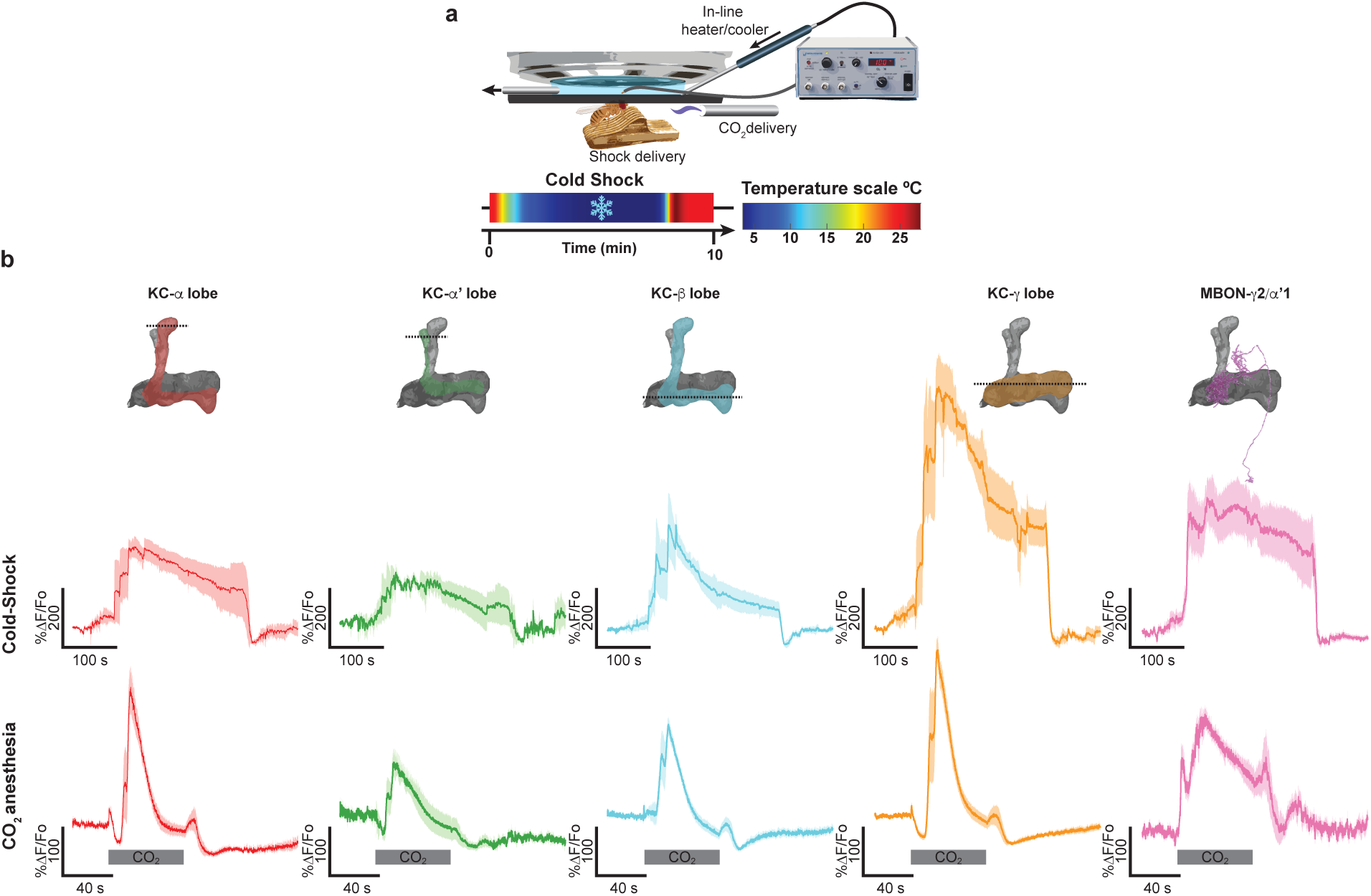
Cold-Shock and CO2 anesthesia induced responses in Kenyon Cells and MBONs. **a,** Diagram of experimental set up to record in vivo anesthesia-induced responses under the microscope. Flies are mounted to a chamber where their body is free, but head if fixed with optical access to their brains. Cold-shock anesthesia is induced by decreasing the temperature from 25°C to 0°C using an in-line heater/cooler Peltier. CO_2_-anesthesia is induced by delivering the gas to the fly antenna through a Teflon tube. The temperature scale shows the change in temperature the fly experiences during the recordings. **b,** KC subtypes and MBON-*g*2*a*’1 calcium traces to cold-shock and CO_2_ induced anesthesia. The onset of CO_2_ anesthesia-induced activity is temporally delayed from the onset of behavioral anesthesia in the fly.

**Extended Data Fig. 6.**
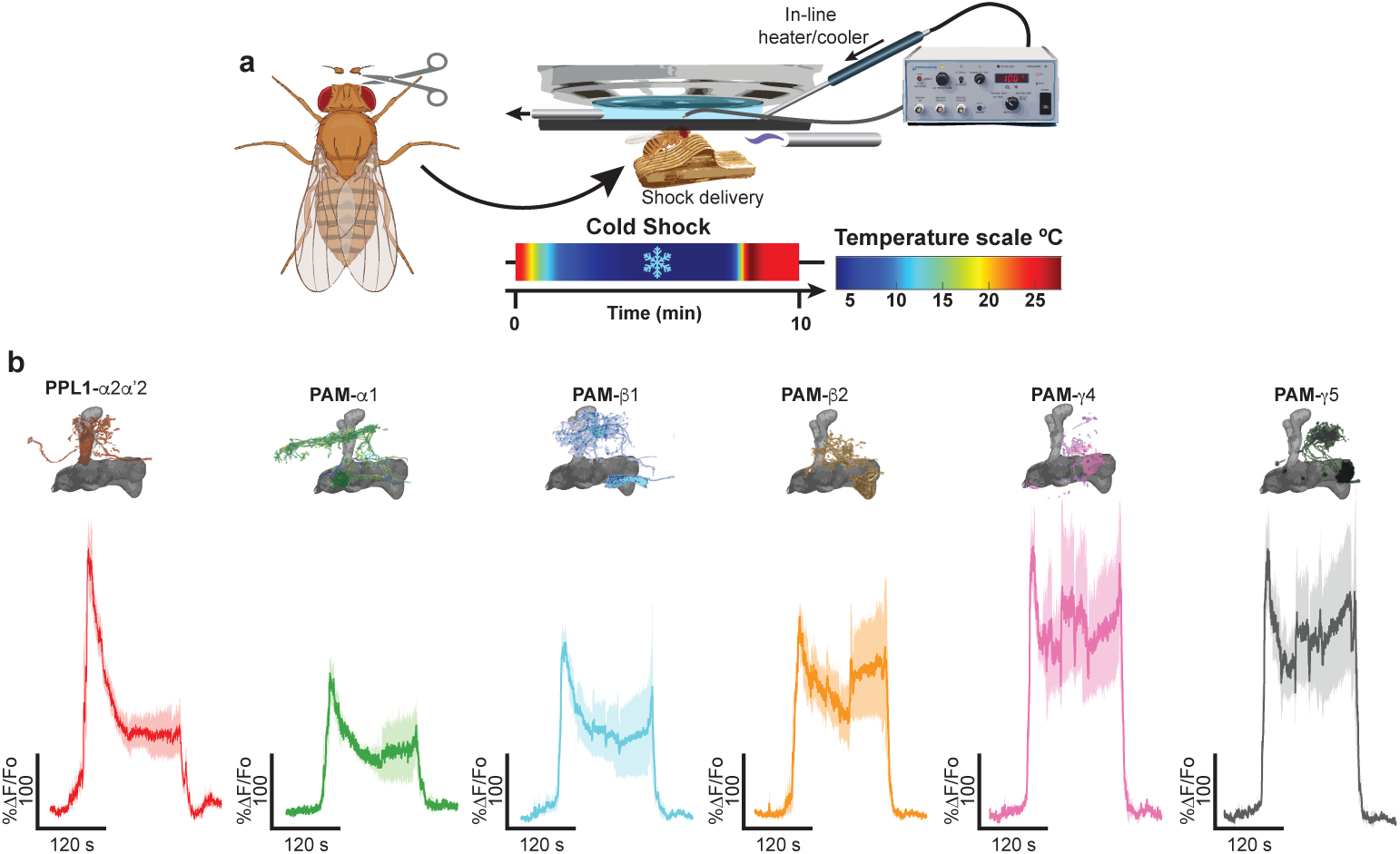
Surgical ablation of antennae does not alter response profile of anesthesia induced activity in dopaminergic neurons. **a,** Diagram of experimental design. The antennae of each fly were surgically ablated and then imaged using the normal cold-shock anesthesia protocol. **b,** Calcium traces for PPL1-a2a’2 and PAM during cold-shock induced anesthesia.

**Extended Data Fig. 7.**
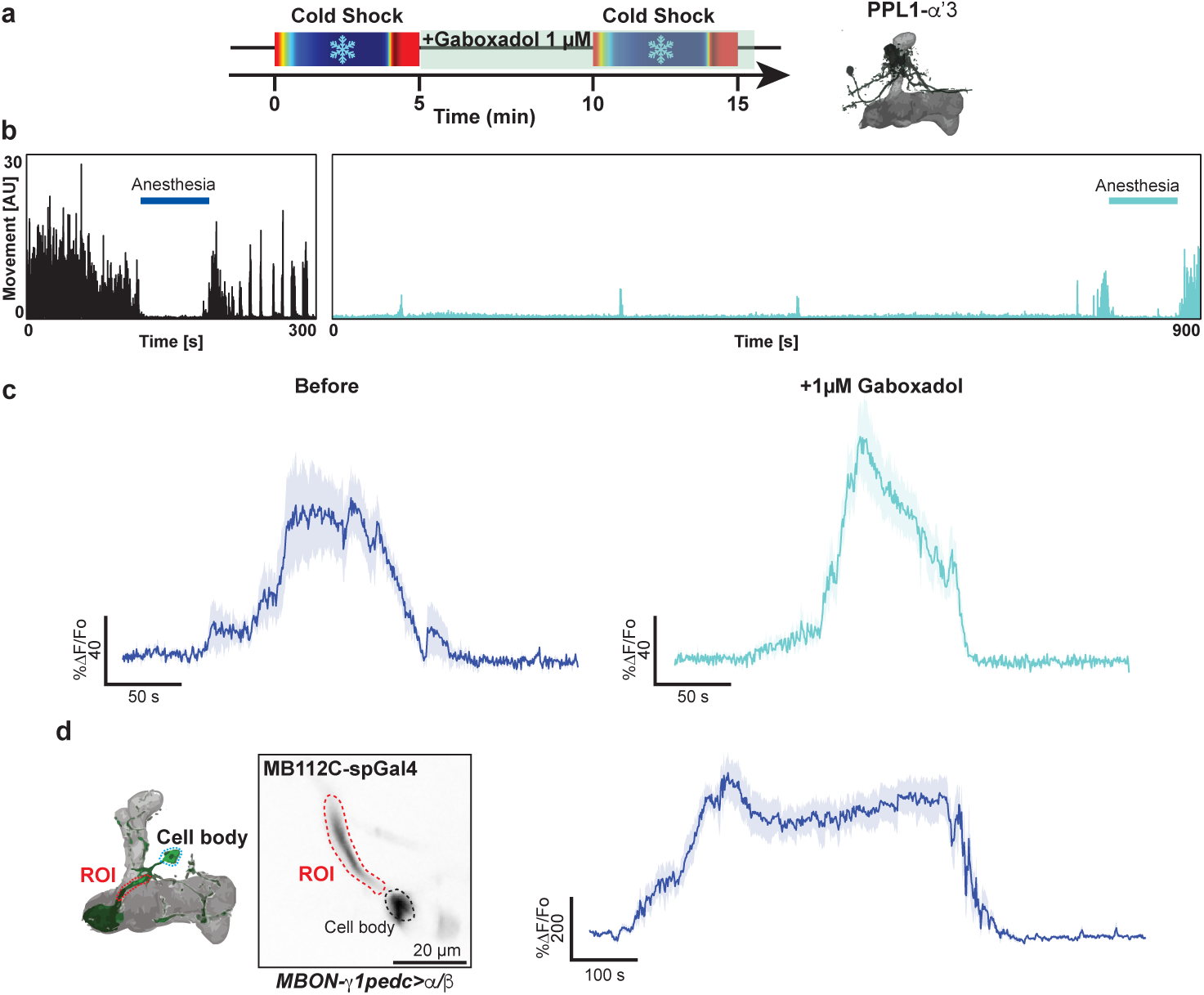
GABA agonist Gaboxadol induces sleep but does not affect cold-shock anesthesia induced response in dopaminergic neurons. **a,** Diagram of experimental design. Each fly was imaged twice for 5 min. Baseline of calcium in PPL1-α’3 was recorded for 10 seconds before cold-shock. Then, saline with 1 mM Gaboxadol is perfused for 10 minutes before a second cold-shock recording was repeated. **b,** Movement of the fly quantified before and during gaboxadol treatment. **c,** Calcium trace before and after gaboxadol treatment, showing no change in anesthesia induced activity. **d,** Mean time series projection of baseline GCaMP6f in MBON-g1pedc>a/b. MBON-g1pedc>a/b GABAergic neuron displays robust calcium activity during cold-shock anesthesia.

**Extended Data Fig. 8.**
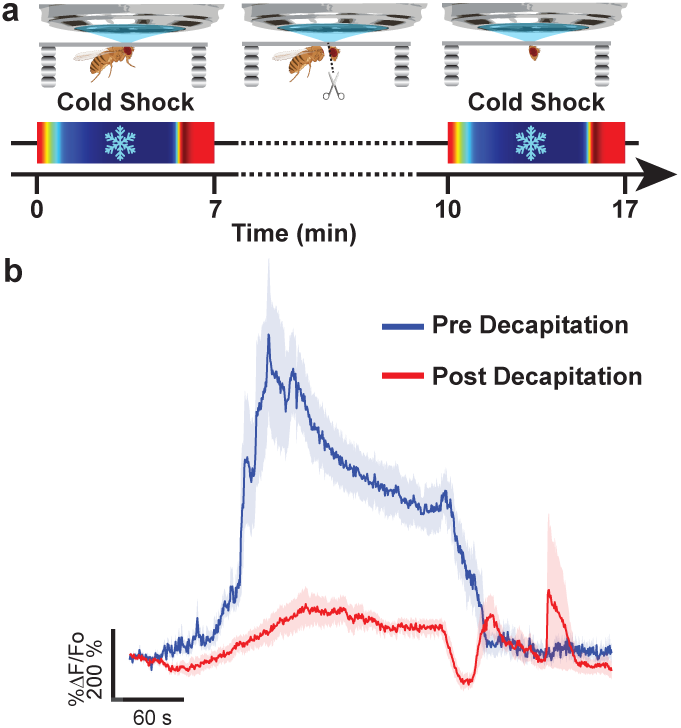
VNC severance abolishes cold-shock induced anesthesia responses in dopaminergic neurons. **a,** Diagram of experimental design. Each fly was imaged twice for 10 min. Baseline of calcium in PPL1-a2a’2 was recorded for 10 s before an initial cold-shock response in intact flies. After the initial recordings, flies were carefully decapitated and a second cold-shock recording was obtained. **b,** Calcium traces showing pre and post decapitation cold-shock anesthesia induced responses.

**Extended Data Fig. 9.**
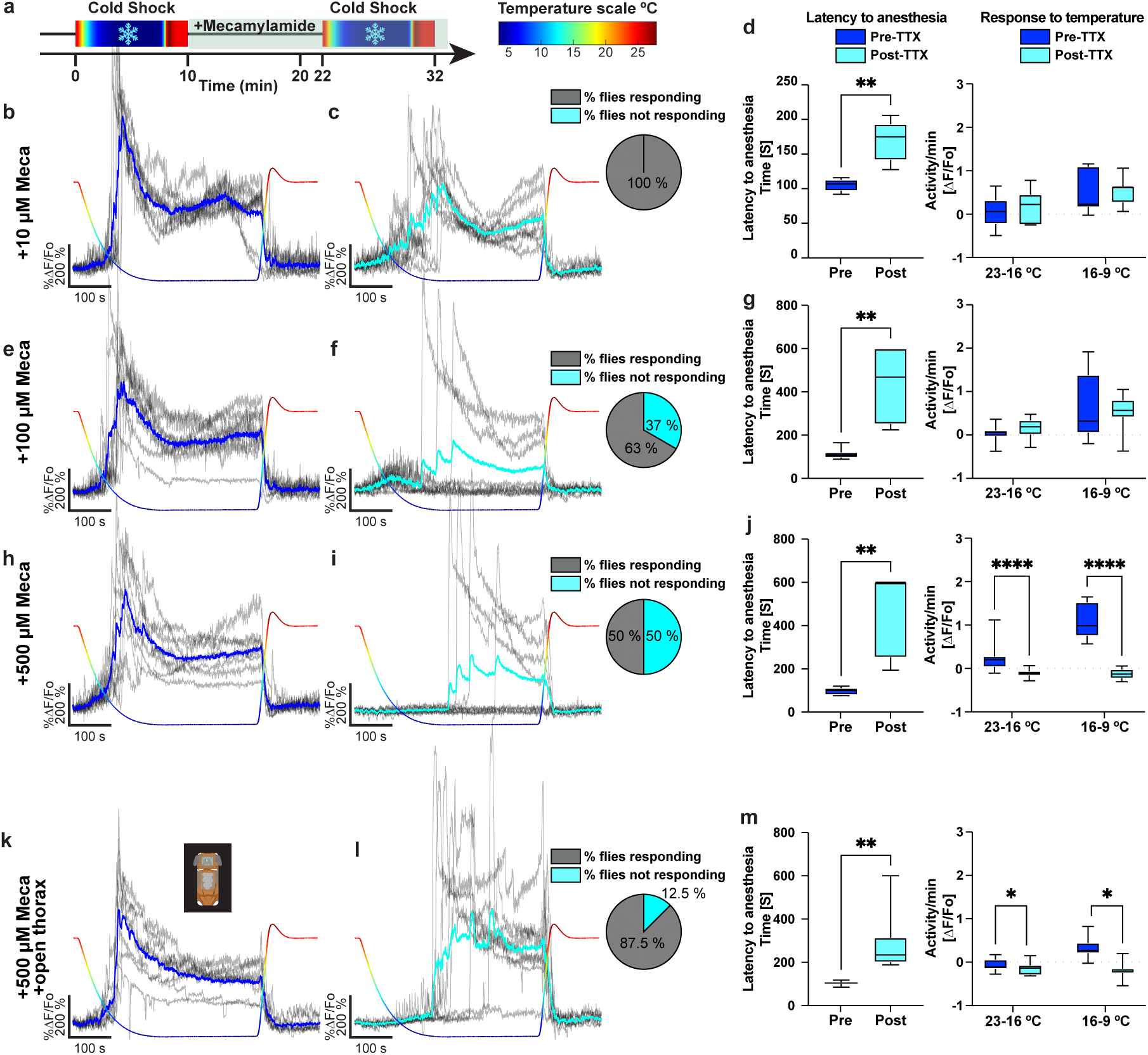
Mecamylamine delays latency to onset of anesthesia-induced response. **a,** Diagram of experimental design. Each fly was imaged twice for 10 min. Baseline of calcium in PPL1-α2α’2 was recorded for 10 s before cold-shock. The temperature returned to 25°C at 7.5 min. Following initial recording, saline containing different concentrations of mecamylamine was perfused for 10 minutes. A second cold-shock recording timeline was repeated post mecamylamine treatment. Temperature recordings are shown as multicolor scale for each calcium trace. Scale is shown with corresponding colors. **b,e,h,k,** Cold-shock induced anesthesia responses in DAN pre mecamylamine perfusion. Gray traces indicate individual fly responses; blue trace is the average anesthesia induced response. **c,** Cold-shock induced anesthesia responses in PPL1 post 10µM mecamylamine perfusion. Pie chart shows that 100% of flies displayed an anesthesia induced response post mecamylamine treatment. **d,** left panel shows latency to onset of anesthesia response pre and post 10 µM mecamylamine treatment. Right panel, calcium response to temperature during two ranges, from 23°C to 16°C and 16°C to 9°C pre and post 10µM mecamylamine treatment. **f,** Cold-shock induced anesthesia responses in PPL1 post 100µM mecamylamine perfusion. Pie chart shows that 37% of flies displayed an anesthesia induced response post mecamylamine treatment. **g,** left panel shows latency to onset of anesthesia response pre and post 100 µM mecamylamine treatment. Right panel, calcium response to temperature during two ranges, from 23°C to 16°C and 16°C to 9°C pre and post 100µM mecamylamine treatment. **i,** Cold-shock induced anesthesia responses in PPL1 post 500µM mecamylamine perfusion. Pie chart shows that 50% of flies displayed an anesthesia induced response post mecamylamine treatment. **j,** left panel shows latency to onset of anesthesia response pre and post 500 µM mecamylamine treatment. Right panel, calcium response to temperature during two ranges, from 23°C to 16°C and 16°C to 9°C pre and post 500µM mecamylamine treatment. **l,** Cold-shock induced anesthesia responses post 500 µM mecamylamine perfusion with VNC access. Pie chart shows that 12.5% flies displayed an anesthesia response post mecamylamine treatment. **m,** left panel shows latency to onset of anesthesia response pre and post 500 µM mecamylamine treatment with VNC access. Right panel, calcium response to temperature during two ranges, from 23°C to 16°C and 16°C to 9°C pre and post 500µM mecamylamine treatment with VNC access. Non-parametric Wilcoxon paired test for pair comparisons. Two-way Anova, for temperature range comparisons followed by Tukey’s multiple comparisons. *p<0.05, **p<0.01, ***p<0.001, ****p<0.0001. Box represents Q1, median, and Q3. Whiskers represents min and max. n=8.

**Extended Data Fig. 10.**
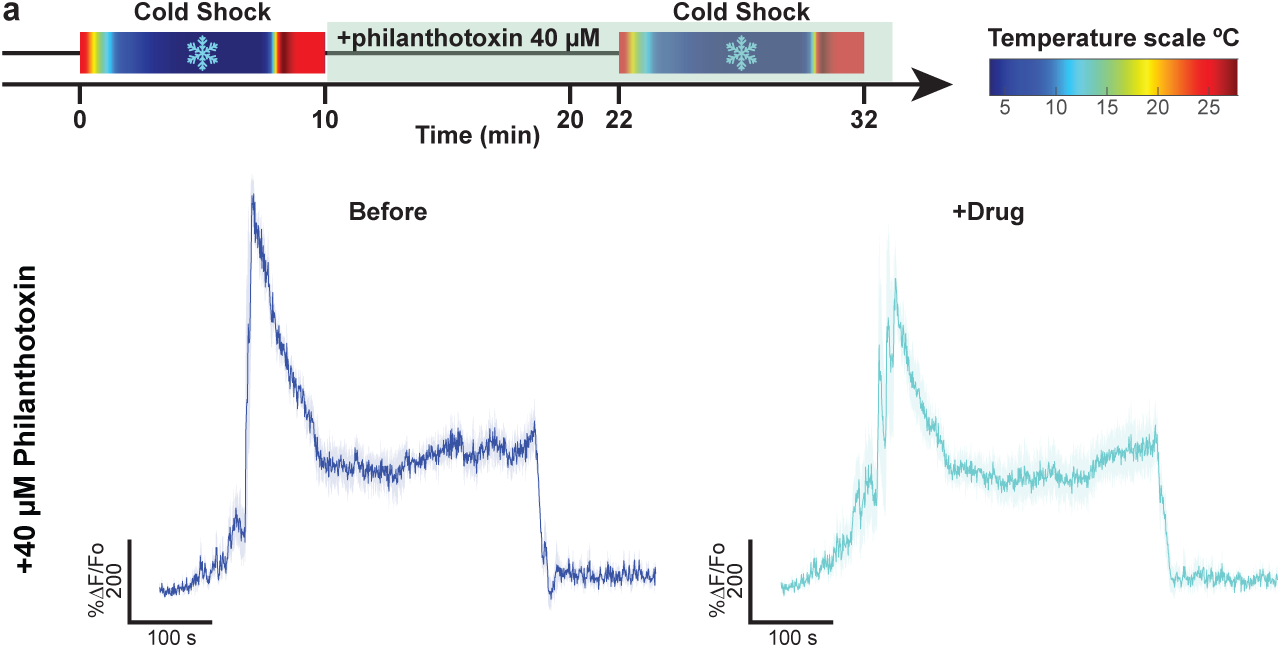
Philanthotoxin does not alter anesthesia induced activity in DANs. **a,** Diagram of experimental design. Each fly was imaged twice for 10 min. Baseline of calcium in PPL1-a2a’2 was recorded for 10 seconds before cold-shock. The temperature was increased back to 25°C at 7.5 min. Then, saline containing 40 µM philanthotoxin was perfused for 10 minutes. A second cold-shock recording timeline was repeated post philanthotoxin treatment. **b,** Cold-shock induced anesthesia responses in PPL1 pre and post 40µM Philanthotoxin perfusion.

**Extended Data Fig. 11.**
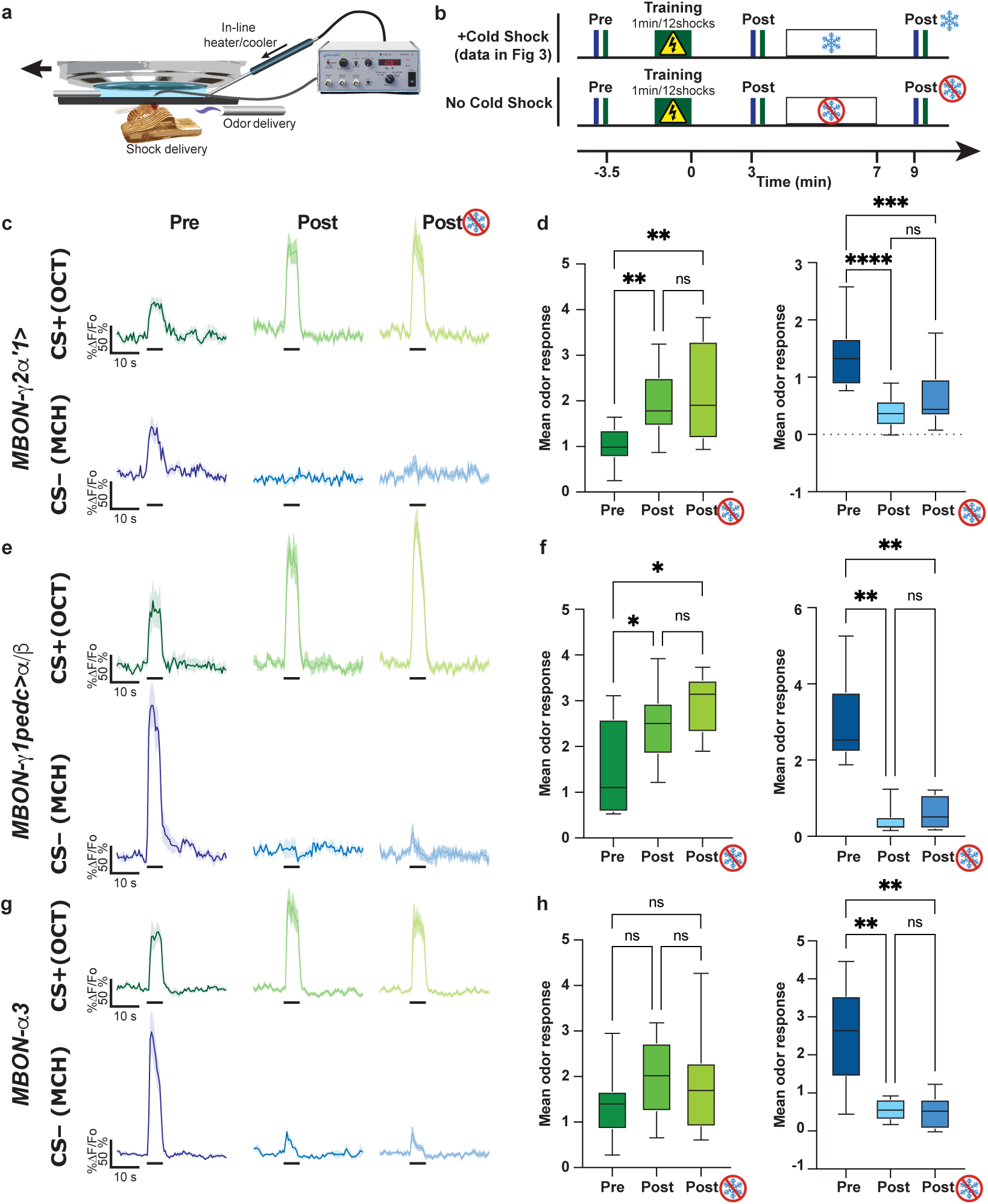
MBON memory traces remain nine minutes post training. **a,** Diagram illustrating in vivo, under the microscope aversive olfactory conditioning training set up and an in-line heater/cooler to deliver cold-shock anesthesia delivery. **b,** Experimental design. Upper panel, Pre-training odor induced calcium response were recorded by presentation of 5-s 4-methylcyclohexanol (MCH) and 3-Octanol (OCT) with a 30-s ISI; 5 minutes later, flies were aversively trained by pairing 1 min presentation of OCT with 12-90 V shocks. Post-training odor induced calcium response were recorded 3 min after. Flies were then subjected to a mock cold-shock under the microscope where saline was perfused but temperature was maintained constant. Two min after, odor responses were recorded. **c,e,g,** Odor response traces to CS+–OCT (top panel) and CS- –MCH (bottom panel) for pre-training, post-training and post-no cold-shock anesthesia for MBON-*γ*2*α*‘1 (**c**), MBON-*γ*1pedc>*α*/*β* (**e**), and MBON-*α*3 (**g**). Thick black bars indicate the timing of odor presentation. **d,** MBON-*γ*2*α*‘1 MCH (CS-) responses were significantly potentiated post-training. OCT (CS+) responses were significantly suppressed post-training. The plasticity to both the CS- and CS+ was maintained following mock cold-shock anesthesia. **f,** MBON-*γ*1pedc>*α*/*β* MCH (CS-) responses were significantly potentiated post training. OCT (CS+) responses were significantly suppressed post-training. Similar to MBON-*γ*2*α*‘1, the plasticity to both the CS-and CS+ was maintained following mock cold-shock anesthesia. **h,** MBON-*α*3 MCH (CS-) responses were significantly potentiated post-training. OCT (CS+) responses were significantly suppressed post-training. Once again, the plasticity to both the CS- and CS+ was maintained following mock cold-shock anesthesia. One-way Anova followed by Tukey’s multiple comparisons test, *p<0.05, **p<0.01, ***p<0.001, ****p<0.0001. Box represents Q1, median, and Q3. Whiskers represents min and max. n=8-12.

**Extended Data Fig. 12.**
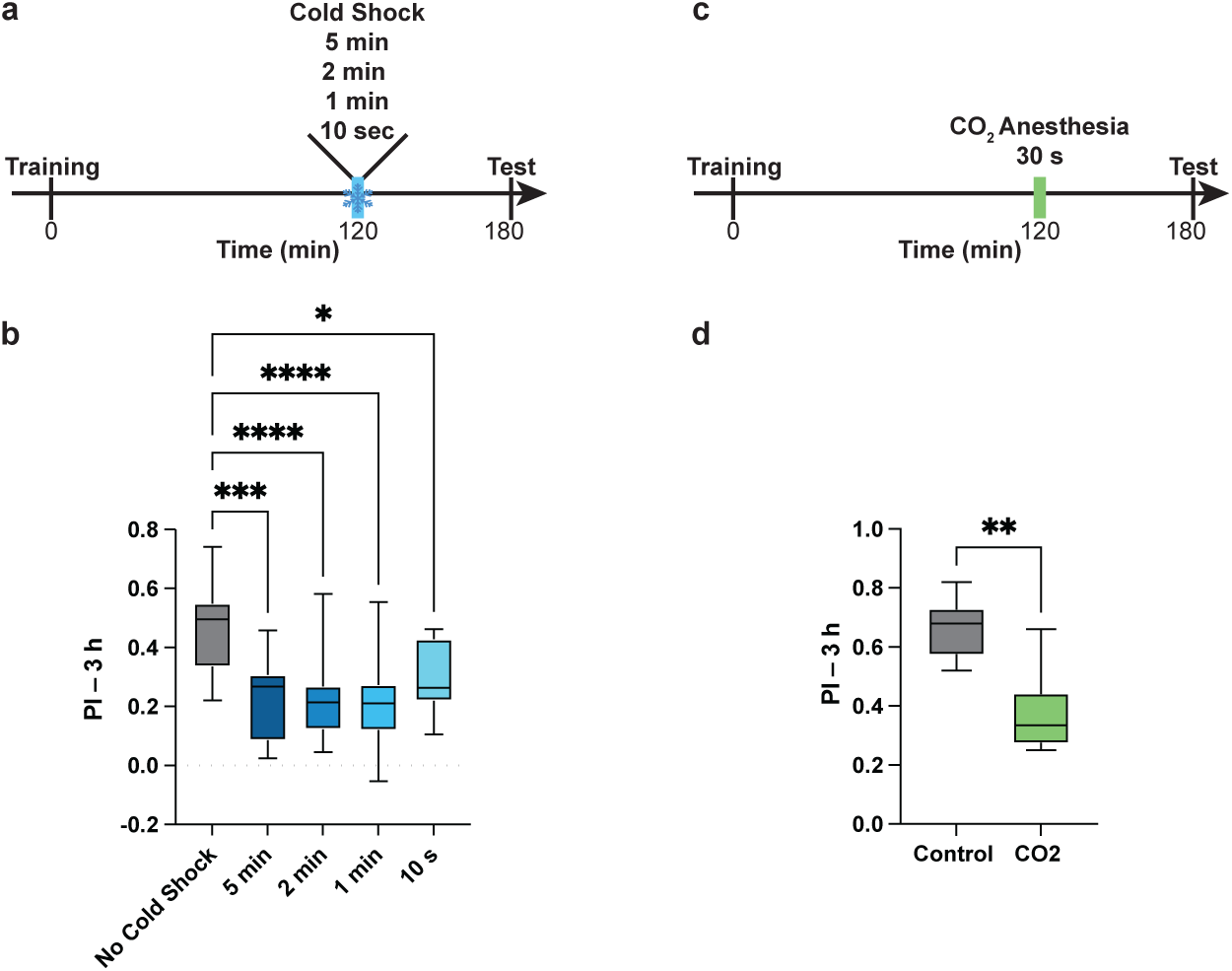
Cold-shock anesthesia as short as ten seconds induces memory loss. **a,** Diagram of experimental protocol. Flies were trained using single cycle aversive olfactory conditioning and cold-shocked for 5, 2, 1 min or 10 s 120 min after training following by a memory test at 180 min post training. **b,** Three-hour memory was significantly impaired in flies subjected to cold-shock anesthesia compared to control animals. **c,** Diagram of experimental protocol. Flies were trained using single cycle aversive olfactory conditioning and given CO_2_ for 30 s at 120 min after training. Flies were subsequently tested for their memory performance at 180 min. **d,** Memory was significantly impaired when flies are CO_2_ anesthetized compared to control animals. One-way Anova followed by Dunnett’s multiple comparisons, *p<0.01, ***p<0.0004, ****p<0.0001. Box represents Q1, median, and Q3. Whiskers represents min and max. n=8-20.

**Extended Data Fig. 13.**
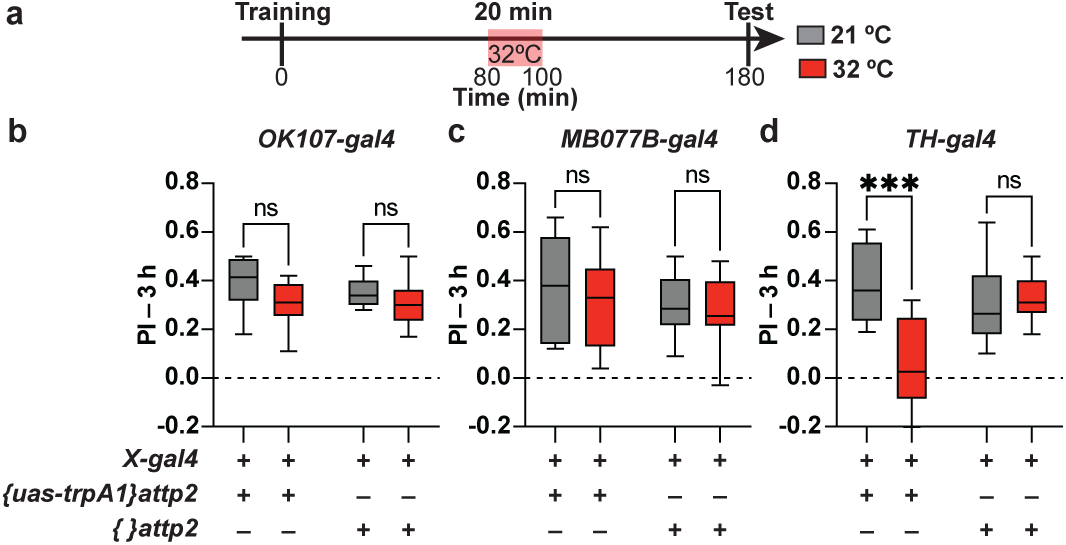
Artificial activation of KCs and MBON-γ2α’1 post-training does not affect memory. **a,** Diagram of experimental protocol. Flies were trained using single cycle aversive olfactory conditioning and placed in 32°C for 20 min, 80 minutes post training. Flies were then tested for memory performance 180 min post-training. **b,** Activation of Kenyon cells does not significantly alter memory performance compared to genetic and temperature-matched control groups. **c,** Activation of MBON-γ2α’1 does not significantly alter memory performance compared to genetic and temperature-matched control groups. **d,** Memory was significantly impaired when DANs (PPL1) were activated compared to genetic and temperature controls. Two-way Anova followed by Fisher’s multiple comparisons, ***p<0.001. Box represents Q1, median, and Q3. Whiskers represents min and max. n=7-8.

## Notes

### Competing Interest Statement

The authors have declared no competing interest.

### Summary of Updates

This version of the manuscript now includes all supplemental data/figures.

## References

1. Berry, J. A., Guhle, D. C. & Davis, R. L. Active forgetting and neuropsychiatric diseases. Molecular Psychiatry 2024 29:9 29, 2810–2820 (2024).

2. Shuai, Y. et al. Forgetting is regulated through Rac activity in Drosophila. Cell 140, 579–589 (2010).

3. Shuai, Y. et al. Dissecting neural pathways for forgetting in Drosophila olfactory aversive memory. Proc Natl Acad Sci U S A 112, E6663–72 (2015).

4. Gao, Y. et al. Genetic dissection of active forgetting in labile and consolidated memories in Drosophila. Proc Natl Acad Sci U S A 10.1073/pnas.1903763116 (2019) doi:10.1073/pnas.1903763116.

5. Berry, J. A., Phan, A. & Davis, R. L. Dopamine Neurons Mediate Learning and Forgetting through Bidirectional Modulation of a Memory Trace. Cell Rep 25, 651–662 e5 (2018).

6. Berry, J., Cervantes-Sandoval, I., Nicholas, E. & Davis, R. Dopamine is required for learning and forgetting in Drosophila. Neuron 74, 530–572 (2012).

7. Berry, J. A., Cervantes-Sandoval, I., Chakraborty, M. & Davis, R. L. Sleep Facilitates Memory by Blocking Dopamine Neuron-Mediated Forgetting. Cell 161, 1656–1667 (2015).

8. Cervantes-Sandoval, I., Chakraborty, M., MacMullen, C. & Davis, R. L. Scribble Scaffolds a Signalosome for Active Forgetting. Neuron 90, 1230–1242 (2016).

9. Cervantes-sandoval, I., Davis, R. L. & Berry, J. A. Rac1 Impairs Forgetting-Induced Cellular Plasticity in Mushroom Body Output Neurons. Front Cell Neurosci 14, 1–11 (2020).

10. Owald, D. & Waddell, S. Olfactory learning skews mushroom body output pathways to steer behavioral choice in Drosophila. Current Opinion in Neurobiology Preprint at 10.1016/j.conb.2015.10.002 (2015).

11. Burke, C. J. et al. Layered reward signalling through octopamine and dopamine in Drosophila. Nature 492, 433–437 (2012).

12. Liu, C. et al. A subset of dopamine neurons signals reward for odour memory in Drosophila. Nature 488, 512–516 (2012).

13. Aso, Y. et al. The neuronal architecture of the mushroom body provides a logic for associative learning. Elife 3, (2014).

14. Aso, Y. et al. Mushroom body output neurons encode valence and guide memory-based action selection in Drosophila. Elife 3, (2014).

15. Hige, T., Aso, Y., Modi, M. N., Rubin, G. M. & Turner, G. C. Heterosynaptic Plasticity Underlies Aversive Olfactory Learning in Drosophila. Neuron 88, 985–998 (2015).

16. Aso, Y. & Rubin, G. M. Dopaminergic neurons write and update memories with cell-type-specific rules. Elife 5, (2016).

17. Martinez-Cervantes, J., Shah, P., Phan, A. & Cervantes-Sandoval, I. High order unimodal olfactory sensory preconditioning in Drosophila. Elife 11, (2022).

18. Séjourné, J. et al. Mushroom body efferent neurons responsible for aversive olfactory memory retrieval in Drosophila. Nat Neurosci 14, 903–910 (2011).

19. Plaçais, P.-Y. Y., Trannoy, S., Friedrich, A. B., Tanimoto, H. & Preat, T. Two pairs of mushroom body efferent neurons are required for appetitive long-term memory retrieval in Drosophila. Cell Rep 5, 769–780 (2013).

20. Pai, T.-P. et al. Drosophila ORB protein in two mushroom body output neurons is necessary for long-term memory formation. Proceedings of the National Academy of Sciences 110, 7898–7903 (2013).

21. Owald, D. et al. Activity of defined mushroom body output neurons underlies learned olfactory behavior in Drosophila. Neuron 86, 417–427 (2015).

22. Perisse, E. et al. Aversive Learning and Appetitive Motivation Toggle Feed-Forward Inhibition in the Drosophila Mushroom Body. Neuron 90, 1086–1099 (2016).

23. Huang, N. et al. Rapid memory shift between different synaptic ensembles promotes forgetting in Drosophila. Current Biology 35, 3119–3132.e4 (2025).

24. Sabandal, J. M., Berry, J. A. & Davis, R. L. Dopamine-based mechanism for transient forgetting. Nature 1–5 (2021) doi:10.1038/s41586-020-03154-y.

25. Handler, A. et al. Distinct Dopamine Receptor Pathways Underlie the Temporal Sensitivity of Associative Learning. Cell 10.1016/j.cell.2019.05.040 (2019) doi:10.1016/j.cell.2019.05.040.

26. Quinn, W. G. & Dudai, Y. Memory phases in Drosophila. Nature 262, 576–577 (1976).

27. Tomchik, S. M. Dopaminergic neurons encode a distributed, asymmetric representation of temperature in Drosophila. J Neurosci 33, 2166 (2013).

28. Bonheur, M. et al. A rapid and bidirectional reporter of neural activity reveals neural correlates of social behaviors in Drosophila. Nat Neurosci 26, 1295–1307 (2023).

29. Cervantes-Sandoval, I., Phan, A., Chakraborty, M. & Davis, R. L. Reciprocal synapses between mushroom body and dopamine neurons form a positive feedback loop required for learning. Elife 6, (2017).

30. Takemura, S. Y. et al. A connectome of a learning and memory center in the adult Drosophila brain. Elife 6, (2017).

31. Stahl, A. et al. Associative learning drives longitudinally graded presynaptic plasticity of neurotransmitter release along axonal compartments. Elife 11, 1–23 (2022).

32. Brown, E. N., Purdon, P. L. & Van Dort, C. J. General Anesthesia and Altered States of Arousal: A Systems Neuroscience Analysis. Annu Rev Neurosci 34, 601 (2011).

33. Adam, E., Kwon, O., Montejo, K. A. & Brown, E. N. Modulatory dynamics mark the transition between anesthetic states of unconsciousness. Proc Natl Acad Sci U S A 120, e2300058120 (2023).

34. Purdon, P. L. et al. Electroencephalogram signatures of loss and recovery of consciousness from propofol. Proc Natl Acad Sci U S A 110, E1142 (2013).

35. Leao, A. A. P. SPREADING DEPRESSION OF ACTIVITY IN THE CEREBRAL CORTEX. 10.1152/jn.1944.7.6.359 7, 359–390 (1944).

36. Connolly, J. B. et al. Associative learning disrupted by impaired Gs signaling in Drosophila mushroom bodies. Science (1979) 274, 2104–2107 (1996).

37. Jenett, A. et al. A GAL4-driver line resource for Drosophila neurobiology. Cell Rep 2, 991–1001 (2012).

38. Chen, T. W. et al. Ultrasensitive fluorescent proteins for imaging neuronal activity. Nature 499, 295–300 (2013).

39. Nern, A., Pfeiffer, B. D., Svoboda, K. & Rubin, G. M. Multiple new site-specific recombinases for use in manipulating animal genomes. Proceedings of the National Academy of Sciences 108, (2011).

40. Bushey, D. et al. RubyACRs Enable Red-Shifted Optogenetic Inhibition in Freely Behaving Drosophila. Elife 14, (2025).

41. Pech, U. et al. Mushroom body miscellanea: transgenic Drosophila strains expressing anatomical and physiological sensor proteins in Kenyon cells. Front Neural Circuits 7, 147 (2013).

42. Shah, P. & Cervantes-Sandoval, I. In Vivo Imaging of Neural Activity in Unanesthetized Drosophila Adult Flies. J Vis Exp 2025-June, (2025).

